# Transcription Factors DksA and PsrA are synergistic contributors to *L. pneumophila* virulence in *Acanthamoeba castellanii* protozoa

**DOI:** 10.1101/2024.07.09.602712

**Authors:** Christopher I. Graham, Andrew J. Gierys, Teassa L. MacMartin, Tiffany V. Penner, Jordan C. Beck, Gerd Prehna, Teresa R. de Kievit, Ann Karen C. Brassinga

## Abstract

The environmental bacterium *Legionella pneumophila*, an intracellular parasite of free-living freshwater protozoa as well as an opportunistic human pathogen, has a biphasic lifestyle. The switch from the vegetative replicative form to the environmentally resilient transmissive phase form is governed by a complex stringent response-based regulatory network that includes RNA polymerase co-factor DksA. Here we report that, through a dysfunctional DksA mutation (DksA1), a synergistic interplay was discovered between DksA and transcription regulator PsrA using the *Acanthamoeba castellanii* protozoan infection model. Surprisingly, *in trans* expression of PsrA partially rescued the growth defect of a *dksA1* strain. While *in trans* expression of DksA expectantly could fully rescued the growth defect of the *dksA1* strain, it could also surprisingly rescue the growth defect of a Δ*psrA* strain. Conversely, the severe intracellular growth defect of a Δ*dksA* strain could be rescued by *in trans* expression of DksA and DksA1, but not PsrA. *In vitro* phenotypic assays show that either DksA or DksA1 were required for extended culturability of bacterial cells, but normal cell morphology and pigmentation required DksA only.

Comparative structural modeling predicts that the DksA1 mutation affects coordination of Mg^2+^ into the active site of RNAP compromising transcription efficiency. Taken together, we propose that PsrA transcriptionally assists DksA in the expression of select transmissive phase traits.

Additionally, *in vitro* evidence suggests that the long chain fatty chain metabolic response is mediated by PsrA together with DksA inferring a novel regulatory link to the stringent response pathway.

## Introduction

*Legionella pneumophila*, a Gram-negative gammaproteobacterium, is commonly found in piped water distribution systems where it is found as an intracellular parasite of free-living bacterivorous protozoa (Rowbowtham, 1986; Steinert *et al.,* 2002; Newton *et al.,* 2010). These environments are not only amenable to bacterial colonization and proliferation, but can also aerosolize *Legionella*-laden water droplets that can be inhaled by susceptible individuals, thereby facilitating bacterial infection of alveolar macrophages resulting in the atypical pneumonia Legionnaires’ disease (Newton *et al.,* 2010; Oliva *et al.,* 2018; Mondino *et al.,* 2020).

*L. pneumophila* is well adapted for its intracellular lifestyle in protozoa, and these adaptations also enable human infection due to highly conserved eukaryotic cellular biology (Newton *et al.,* 2010; Mondino *et al.,* 2020). A core driver of infection is the Dot/Icm Type IV secretion system (T4SS), an apparatus that enables secretion of a diverse repertoire of ≥ 330 effectors into the host cytoplasm that together act to modify host cellular physiology upon uptake of the bacteria (Finsel and Hilbi, 2015; Burstein *et al*., 2016; Gomez-Valero *et al.,* 2019). Such modifications include subversion of phagosome trafficking to prevent fusion of the bacteria-containing phagosome with lysosomes, and conversion of the phagosome into a replicative niche referred to as the *Legionella*-containing vacuole (LCV) (Tilney *et al.,* 2001; Kagan and Roy, 2002; Isberg *et al*., 2009). As infection progresses within a protozoan host cell, *L. pneumophila* exhibits a biphasic lifecycle, alternating between a vegetative replicative phase (RP) forms found primarily in the LCV, and transmissive phase (TP) forms that develop upon depletion of host-derived nutrients in anticipation of bacterial release to the environment. The physiology and ultrastructure of TP cells is optimized for environmental survival and resistance to stress, with expressed virulence factors enabling infection of a new host (Molofsky and Swanson, 2004; Robertson *et al*., 2014; Oliva *et al.,* 2018).

The biphasic switch between RP and TP is controlled by a stringent response-based regulatory network activated by accumulation of alarmone guanosine tetraphosphate (ppGpp), which is synthesized by RelA and SpoT in response to nutritional deprivation. Synthase RelA senses low amino acid concentrations (Zusman *et al.,* 2002), whereas bifunctional synthase and hydrolase SpoT senses fatty acid flux via acyl carrier proteins (ACP) (Edwards *et al.,* 2009; Dalebroux *et al.,* 2010). Accumulated ppGpp is associated with activation of the LetA/LetS two component system (TCS), which in turn induces transcription of non-coding RNAs (ncRNAs) RsmY and RsmZ (Hammer *et al*., 2002; Rasis and Segal, 2009; Sahr *et al.,* 2009; Nevo *et al.,* 2014). These ncRNAs target and sequester the RNA-binding protein CsrA, thus modulating CsrA-mediated post-transcriptional regulation of transcripts associated with the transition to TP. Such transcripts include those encoding Dot/Icm effectors and those linked to cellular metabolism (Häuslein *et al.,* 2017; Sahr *et al*., 2017). Other TCS also play a pivotal role in bacterial differentiation and are often required for virulence in host cells, such as CpxR/A (Feldheim *et al.,* 2016; Tanner *et al.,* 2016); PmrA/B (Zusman *et al.,* 2007; Al-Khodor *et al.,* 2009) and quorum sensing via LqsS/T/R (Tiaden *et al.,* 2007; Tiaden *et al.,* 2008; Schell *et al.,* 2014; Personnic *et al.,* 2018). In addition, a subset of transcriptional regulators plays an important role in TP induction. IHF assists LetA in the upregulation of RsmY and RsmZ, and is required for full cyst-like form differentiation and virulence in *A. castellanii* (Morash *et al*., 2009; Pitre *et al*., 2013). OxyR regulates transcription of the operon encoding the CpxR/A TCS, linking it to virulence in *A. castellanii* (Tanner *et al.,* 2017). Recently, the autorepressor PsrA was found to be required for optimal intracellular growth and cyst-like form biogenesis in *A. castellanii* (Graham *et al*., 2021). While the genetic targets of PsrA have not yet been identified, a potential linkage between PsrA and the core stringent response network has been described with focus on fatty acid flux and an associated ACP (Graham *et al.,* 2021).

Accumulated ppGpp also directly affects transcriptomic changes associated with the transition from RP to TP. For example, expression of stationary phase sigma factor RpoS is increased (Dalebroux *et al*., 2010), which is required for the expression of TP traits and intracellular growth in protozoa such as *A. castellanii* (Hales and Shuman, 1999; Bachman and Swanson, 2001; Zusman *et al.,* 2002; Abu Zant *et al.,* 2006; Hovel-Miner *et al.,* 2009). While the precise mechanism has not yet been ascertained in *L. pneumophila*, ppGpp influences expression most likely by binding directly to RNA polymerase (RNAP), as has been described in *E. coli* where ppGpp binds with equal affinity either to a site encompassed by the ω and β’ subunits, or at the interface between RNAP and secondary channel co-factor DksA. DksA features a pair of conserved aspartate residues to enable ppGpp binding (Perederina *et al.,* 2004; Ross *et al*., 2013; Ross *et al*., 2016; Myers *et al*., 2020). The latter event results in a conformational change in both ppGpp and DksA affecting the stability of the open complex generated by the RNAP holoenzyme thereby altering the efficiency of transcription initiation (Chatterji *et al*., 1998; Lemke *et al*., 2011; Gummesson *et al.,* 2013; Ross *et al*., 2016; Molodtsov *et al*., 2018; Travis and Schumacher, 2021). DksA can act independently of ppGpp, compensate for its absence if overexpressed, or even act antagonistically to ppGpp at different promoters (Magnussun *et al.,* 2007; Aberg *et al.,* 2008; Potrykus and Cashel, 2008; Aberg *et al*., 2009). DksA exists in a dynamic equilibrium with structural homologs GreA and GreB in *E. coli*, which compete with DksA for secondary channel binding. If either GreA or GreB binds, then DksA-independent ppGpp binding to RNAP still occurs to influence expression (Vinella *et al.,* 2012).

*L. pneumophila* encodes both GreA and DksA orthologs, although GreA has not yet been targeted for study. Like its *E. coli* ortholog, DksA acts independently of, or in coordination with, ppGpp to regulate genes encoding TP traits (Dalebroux *et al.,* 2010). Previously, DksA was shown to be required in a ppGpp-dependent manner for TP differentiation including flagellar biosynthesis. Likewise, DksA and ppGpp are both required for optimal intracellular growth in *A. castellanii*. Conversely, DksA, but not ppGpp, is dispensable for intracellular growth in murine macrophages, indicating that ppGpp may act independently of DksA in *L. pneumophila* (Dalebroux *et al.,* 2010). Thus, the contributions of ppGpp and DksA to the stringent response, are well characterized among the proteobacteria though the mechanisms and targets may vary between them (Gourse *et al*., 2018).

In this study, we describe an interplay between DksA and PsrA in the regulation of virulence in protozoa, discovered through a hypomorphic mutant of DksA (DksA1). We show that nominal intracellular growth can be fully achieved with DksA compensating for lack of PsrA, but only partially achieved with PsrA supplementing DksA1, suggesting that PsrA transcriptionally assists DksA. Structural modelling of DksA1 suggests that the transcription efficiency of this RNAP co-factor is compromised, thereby highlighting the proposed synergistic role of PsrA. Further, only DksA is essential for the *in vitro* phenotypic traits of extended culturability, cell morphology and pigmentation. Conversely, *in vitro* evidence suggests that the long chain fatty chain metabolic response is mediated by PsrA together with DksA. Taken together, we show that PsrA and DksA have distinct and overlapping biological roles in the regulation of TP highlighting a complex regulatory interaction between these two regulatory factors.

## Methods

### Structural Predictions and Modelling

The amino acid sequences of DksA and DskA1 were submitted to the collabfold2 sever (Pettersen *et al*. 2021) with default settings. Top rank #1 models were selected for modelling and analysis in UCSF ChimeraX (Pettersen *et al*., 2021). Structural overlays between Lp02 DksA, DksA1, and DksA (5VSW) (Molodtsov *et al*., 2018) were aligned with Matchmaker. Conservation alignments were performed using Consurf (Landau *et al*., 2005; Ashkenazy *et al*., 2016) with the Alphafold2 model of Lp02 DksA.

### Culture of E. coli and L pneumophila

*Legionella pneumophila* Philadelphia-1 derivative Lp02 parental and isogenic strains, as described in Table 1, were grown on Buffered Charcoal Yeast Extract (BCYE) agar plates (Feeley *et al*., 1979) at 37°C and 5% CO_2_, or in Buffered-Yeast Extract (BYE) broth in tubes on a roller drum at 37°C, supplemented as required with thymidine (100 μg/mL), sucrose (5% w/v) or kanamycin sulphate (25 μg/mL). *E. coli* was utilized for cloning and plasmid propagation purposes [DH5α for *in trans* expression vector pJB908 or pThy (pBH6119), and DH5α λpir (suicide vector pSR47S) was grown on LB agar plates, or broth in tubes on a roller drum, supplemented as required with ampicillin (100 μg/mL) or kanamycin sulphate (40 μg/mL) and incubated at 37°C.

**Table 1.** List of strains and plasmids

| Number | Name | Description | Source/reference |
| --- | --- | --- | --- |
| ***E. coli*** |  |  |  |
|  | DH5α | F' *endA1 hsdR17(r<sub>k</sub>- m<sub>k</sub>-) supE44 thi-1 recA1 gyrA (Nal<sup>r</sup>) relA1 Δ(lacZYA-argF)U169 deoR(φ80lacΔ(lacZ)M15)* | New England Biolabs |
|  | DH5α λpir | K-12 F- φ80lacZΔM15 *endA recA hsdR17 (rm-mK+)* *supE44 thi-1 gyrA96 relA1 Δ(lacZYA-argF) U169 λpir* | M. Swanson (Bryan *et al.*, 2013) |
| ***L. pneumophila*** |  |  |  |
| C858 | Lp02 | Str<sup>r</sup>, Thy<sup>-</sup>, HsdR<sup>-</sup> derivative of Philadelphia-1 strain | M. Swanson (Berger and Isberg, 1993) |
| KB265 | Lp03 | Lp02 Δ*dotA* mutant | R. Isberg (Berger and Isberg, 1993) |
| CG859 | Δ*psrA* | Lp02 Δ*psrA* (unmarked, in frame deletion mutant) | Graham *et al.*, (2021) |
| CG860 | Lp02 pThy | Lp02 pBH6119 | Tanner *et al.*, (2016) |
| JT708 | Δ*dotA* pThy | Lp03 pBH6119 | Tanner *et al.*, (2016) |
| CG861 | Lp02 pVect-*psrA* | Lp02 pJB908-*psrA* | Graham *et al.*, (2021) |
| CG862 | Δ*psrA* pThy | Δ*psrA* pThy | Graham *et al.*, (2021) |
| CG863 | Δ*psrA* pVect-*psrA* | Δ*psrA* pJB908-*psrA* | Graham *et al.*, (2021) |
| CG799 | *dksA1* | Lp02 *dksA1* (*dksA1*[Δ170-194]); spontaneous 24 bp in-frame deletion | This study |
| CG819 | *dksA1* pThy | *dksA1* pBH6119 | This study |
| CG813 | *dksA1*Δ*psrA* | Lp02 *dksA1* Δ*psrA* | This study |
| CG826 | *dksA1* Δ*psrA* pThy | *dksA1* Δ*psrA* pBH6119 | This study |
| CG1053 | Δ*dksA* | Lp02 Δ*dksA* (unmarked, in frame deletion mutant of *dksA*) | This study |
| CG1054 | Δ*dksA* Δ*psrA* | Δ*dksA* Δ*psrA* | This study |
| CG1055 | Δ*dksA* pThy | Δ*dksA* pBH6119 | This study |
| CG1056 | Δ*dksA* Δ*psrA* pThy | Δ*dksA* Δ*psrA* pBH6119 | This study |
| CG1057 | Δ*dksA* pVect-*dksA* | Δ*dksA* pBH6119 | This study |
| CG1059 | Lp02 pVect-*dksA* | Lp02 pJB908-*dksA* | This study |
| CG1060 | Δ*psrA* pVect-*dksA* | Δ*psrA* pJB908-*dksA* | This study |
| CG1061 | Δ*dksA* pVect-*psrA* | Δ*dksA* pJB908-*psrA* | This study |
| CG1101 | Δ*dksA* pVect-*dksA1* | Δ*dksA* pJB908-*dksA1* | This study |
| CG1102 | Lp02 pVect-*dksA1* | Lp02 pVect-*dksA1* | This study |
| CG1146 | *dksA1* pVect-*dksA* | *dksA1* pVect-*dksA* | This study |
| **Plasmids** |  |  |  |
|  | pJB908 | RSF1010 cloning vector, *tdΔi*, Amp<sup>r</sup> Thy<sup>+</sup> | J. Vogel (Sexton *et al.*, 2004) |
|  | pBH6119 | RSF1010 promoterless GFP vector; Amp<sup>r</sup>, Thy<sup>+</sup> | M. Swanson (Hammer *et al.*, 2002) |
|  | pSR47S | oriTRP4 oriR6K *kan<sup>r</sup> sacB* | J. Vogel (Merriam *et al.*, 1997) |
| pPP619 | pVect-*psrA* | pJB908-*psrA* | Graham *et al.*, (2021) |
| pCG1051 | pVect-*dksA* | pJB908-*dksA* | This study |
| pCG1080 | pVect-*dksA1* | pJB908-*dksA1* allele recovered from spontaneous<br>*dksA1* Lp02 strain CG799 | This study |
| pCG1069 | pSR47S-5'flk-Δ*dksA*-<br>3'flk | pSR47S with 5' and 3' flanking regions for Δ*dksA* | This study |

### Molecular methods/reagents, cloning, and sequencing

Chemicals were procured form VWR (Ottawa, ON, Canada), Thermo-Fisher (Mississauga, ON, Canada), or Sigma-Aldrich (Oakville, ON, Canada). Restriction endonuclease and PCR polymerase (Taq, Q5), and other cloning reagents were procured from New England Biolabs (Whitby, ON, Canada). Oligonucleotide primers, ordered from Life Technologies (Thermo-Fisher), were utilized under conditions listed in Table 2. Nucleic acid purification kits were purchased from Qiagen (Toronto, ON, Canada). Sanger sequencing for routine verification of constructs and mutant validations was done on a fee-based service by The Centre for Applied Genomics (The Hospital for Sick Children, Toronto, ON, Canada). Whole genome sequencing was undertaken by phenol-chloroform isolation of genomic DNA followed by sequencing via Pacbio SMRT at Genome-Québec (Montreal, QC, Canada) and assembled using Geneious^TM^ software.

**Table 2.**
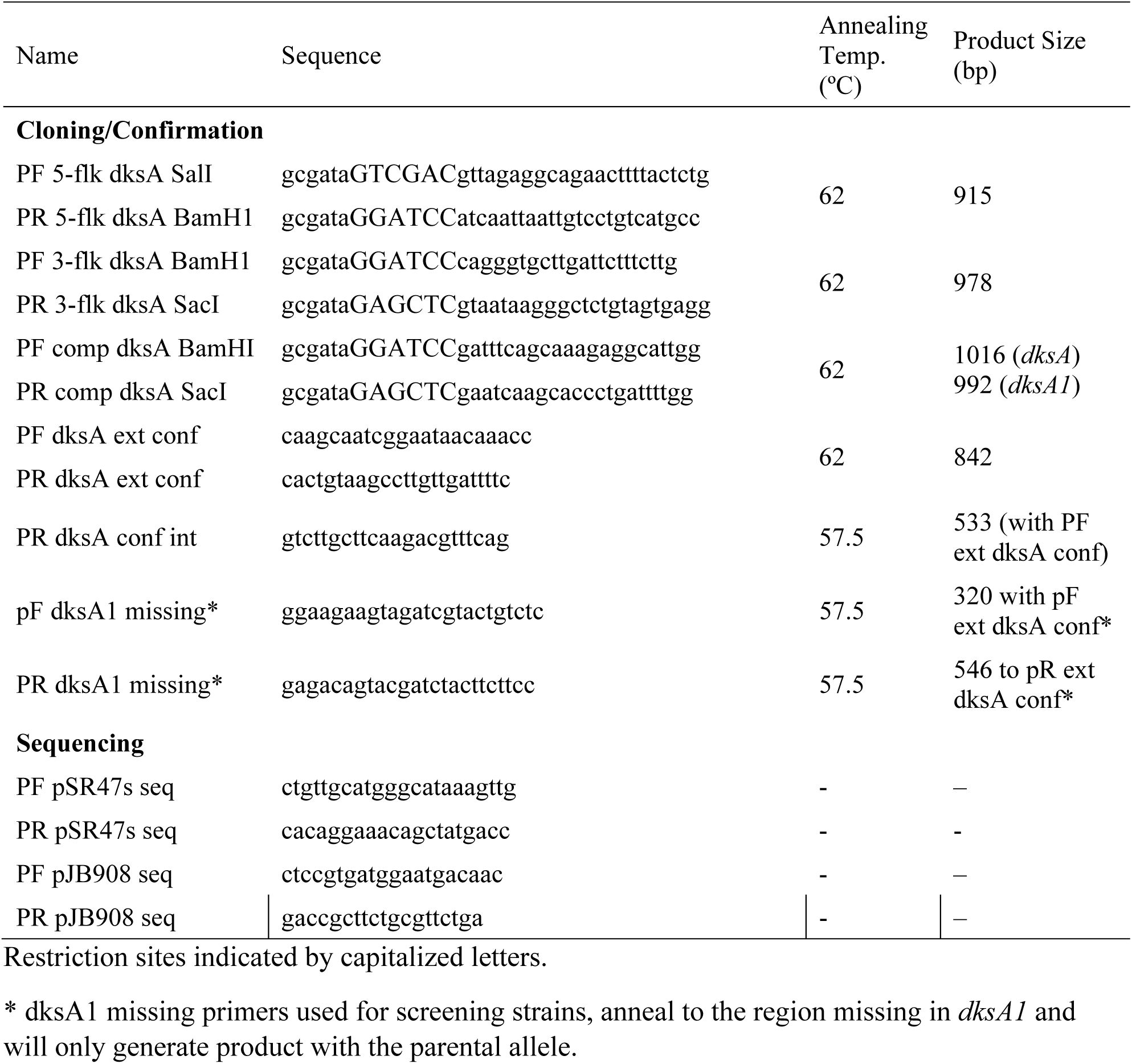
List of oligonucleotides.

### *dskA1* and Δ*dksA* mutant strains

The *dksA1* mutation was identified through whole genome sequencing of a Δ*psrA* mutant strain (i.e. *dksA1* Δ*psrA*) during experiments for Graham *et al*. (2021). The *dskA1* mutation was then traced back to the electrocompetent cell stock to attain the *dksA1* strain via PCR screening using primer set (PF dksA1 missing / PR dksA1 missing) that annealed to the missing 24 bp region within *dksA1* in combination with primer set (PF dksA ext conf / PR dksA ext conf). For these reactions, PF dksA1 missing was paired with PR dksA ext conf, and PF dksA ext conf was paired with PR dksA1 missing, such that each primer set only generated product to identify the *dksA1* allele. For generation of an in-frame Δ*dksA* mutant strain, ∼1000 bp 5′ (primer set PF 5-flk dksA SalI / PR 5-flk dksA BamH1) and 3′ (PF 3-flk dksA BamH1 / PR 3-flk dksA SacI) regions flanking *dksA* were PCR-amplified and cloned into suicide vector pSR47S to use for routine allelic exchange. For generation of *in trans* expression constructs, either *dksA* or *dksA1* were PCR-amplified from Lp02 or *dksA1* mutant CG799, respectively, using primer set (PF comp dksA BamHI / PR comp dksA SacI) and cloned into pJB908, and introduced into parental or isogenic mutant strains by electroporation. The *in trans psrA* expression vector was from a previous study (originally referred to as pComp henceforth renamed to pVect-*psrA* in this study) (Graham *et al*., 2021). Use of pThy empty vector pBH6119 is used to rescue thymidine auxotrophy of Lp02. Plasmid constructs and mutations were routinely verified by sequencing isolated plasmids or PCR amplicon products, and mutant strains were routinely subjected to whole genome sequencing to rule out secondary mutations.

### *In vitro* growth kinetics

*L. pneumophila* strains were grown on plates for three days under standard conditions. Bacterial cell mass was collected with a disposable loop and suspended in pre-warmed BYE broth and normalized to OD_600_ = 0.15. Subsequently 150 µL of this suspension was loaded in triplicate into wells within a 96-well flat bottomed microtitre plate, the lid was sealed with Parafilm, and loaded into a BioTek Synergy HTX plate reader, reading OD_600_ hourly for 24 h at 37°C or every 2 h for 48 h at 25°C with 150 rpm linear shaking. Results were plotted in GraphPad Prism^TM^ 9.1 and analyzed via two-way ANOVA analysis.

### Culture and intracellular kinetics in protozoan *A. castellanii*

*Acanthamoeba castellanii* (ATCC 30234) were propagated in PYG media at 25°C as previously described (Tanner *et al.,* 2016). Briefly, infections were undertaken in AC buffer by placing ∼10^6^ protozoan trophozoites in each well of a set of tissue culture-treated 24-well plates and allowing them to adhere, then overlaying them with bacteria normalized by OD_600_ to a MOI of 0.1 in AC buffer. After 1 h incubation at 25°C, the wells were washed 3 x 1 mL AC buffer, before incubating in 1 mL/well of AC buffer at 25°C until the indicated timepoints. Protozoa were lysed mechanically via vortexing, and the lysate serially diluted in AC buffer onto BCYE-agar plates to enumerate titres. To enhance enumeration of difficult-to-recover avirulent strains, a portion of the lysate was tenfold concentrated by centrifugation at 21,000 x *g* in a microcentrifuge for 2 min, then 0.9 vol of the supernatant drawn off, the pellet resuspended in the remaining volume and spotted alongside the serial dilutions. After counting titres, graphs were plotted and analyzed (two-way ANOVA) in GraphPad Prism^TM^ 9.1.

### Culture and intracellular kinetics in U937 human macrophages

Human U937 monocytic cells were propagated and differentiated into macrophages using phorbol myristate acetate (PMA, Sigma-Aldrich) as previously described (Tanner *et al.,* 2016). Cells were grown in RPMI-1640 media (HyClone, Thermo-Fisher) supplemented with 10% heat inactivated fetal bovine serum (Seradigm, Mississauga, ON) (RPMI + FBS) at 37°C and 5% CO_2_ that was used for all washes and incubations in a pre-warmed state. For infection, ∼ 10^6^ activated U937-derived macrophages were placed into each well of a tissue culture-treated 24 well plate, allowed to adhere, and overlaid with *L. pneumophila* strains normalized by OD_600_ to a MOI of 2, for 1 h.

The bacteria were washed away with 3 x 1 mL washes in RPMI + FBS, followed by an incubation for 1 h with RPMI + FBS with 100 μg/mL gentamicin. After a further 3 x 1 mL washes in RPMI + FBS, the cells were incubated in 1 mL RPMI + FBS until lysis at indicated timepoints. Cells were lysed osmotically by replacing the RPMI + FBS with ice-cold HyClone ultrapure water and repeated pipetting (Thermo-Fisher), then serially diluting the lysate in additional HyClone water and enumerating *L. pneumophila* titres by spotting the dilutions on BCYE-agar plates. After counting, graphs were plotted and analyzed via two-way ANOVA in GraphPad Prism^TM^ 9.1.

### Pigment, morphology and culturability assays

Suspensions of *L. pneumophila* were prepared as per *in vitro* growth kinetics. Pigment assays were undertaken in accordance to Wiater et al. (1994). Briefly, inoculated 5 mL BYE broth cultures were incubated for 72 h. To quantify pigment at 24 h timepoints, 1 mL of culture was withdrawn, pelleted at 21,000 x g in a microcentrifuge for 2 min, after which 150 µL of supernatant was loaded in triplicate into 96 well plates and read for absorbance at 550 nm on a BioTek Synergy HTX plate reader (Wiater *et al*., 1994). After removal of residual supernatant, the pellet was suspended in 1 mL of 1 x PBS pH 7.0 and similarly loaded (150 µL) into a 96 well plate then read for OD_600_. Blank controls were the same volume of BYE or 1 x PBS as appropriate. Results were plotted and analyzed using GraphPad Prism^TM^ (9.1). Photographed samples were prepared similarly and at specified allowed to sit undisturbed for 24 h at ambient temperatures (∼22°C) for cells to sediment out by gravity. Long-term culturability was assessed by inoculating 96 well plates with six wells for each strain, containing 150 µL of an OD_600_ = 0.15 suspension in BYE. The outermost rows and columns were filled with sterile water to reduce evaporation, the plates were sealed with Parafilm and incubated on a shaking platform at 37°C for four days. At indicated intervals, cell aggregates were dispersed with pipetting, the OD_600_ was read with a BioTek Synergy HTX plate reader across the wells containing unharvested culture, and one well of each strain harvested and serially diluted for titre on BCYE agar plates. Results were plotted and analyzed using GraphPad Prism (9.1). For colony-spot morphology, 10 µL of the residual OD_600_ = 0.15 suspension used to prepare these assays was spotted on a BCYE plate with spots spread out to twice the spacing of a standard multi-channel pipette then incubated for 4 days at 37°C and 5% CO_2_, then photographed.

### LCFA assays

Suspensions of *L. pneumophila* strains were prepared as per the *in vitro* kinetics, at OD_600_ = 0.15 in BYE broth, except that media was supplemented with two-fold serially diluted concentrations of palmitic acid (PA) as a 40 mM stock dissolved in 100% ethanol. All broth including PA dilutions, controls and blanks was normalized to 0.5% ethanol. The suspension was then subject to a standard in vitro growth curve as described in the *in vitro* kinetics section.

Inhibition was defined as growth of <50% in comparison to that of the same strain in a BYE + 0.5% ethanol-only control at the end of 24 h. Data were graphed using GraphPad Prism^TM^ 9.1.

### Microscopy

Microscopy and cell lengths were conducted as per Graham *et al.,* (2021). Briefly, broth cultures in BYE broth were inoculated at OD_600_ = ∼0.02 and incubated for 18 h to OD_600_ = 0.5-1.0 at 37°C to enter exponential phase, then were mounted on a 2% w/v agarose pad prepared in water and imaged on an Axio Observer Z1 inverted microscope (Zeiss) equipped with a glycerol-immersion 150X objective. Stationary phase cells were imaged by returning the cultures to the roller drum for a further ∼24 h to achieve stationary phase and imaged similarly. Cell lengths were quantified using the length tool in ImageJ and lengths plotted and analyzed in GraphPad Prism^TM^ (9.1).

## Results

### *dksA1* mutation

In a previous study, we deleted *psrA* from the parental Lp02 strain (Graham *et al*., 2021). Throughout this process, we noted a propensity for some of the isogenic Δ*psrA* strains to acquire secondary mutations in *letS* or *dksA* as discovered through routine whole genome sequencing. One particular Δ*psrA* strain was found to contain an in-frame 24 bp deletion within the *dksA* coding sequence, that resulted in the loss of eight amino acid residues after the 57^th^ residue in the DksA polypeptide (Figure S1A, B). The 57^th^ codon was additionally altered from encoding leucine to phenylalanine. Thus, this allele was denoted as the *dksA1* mutation that is characterized in this study.

As the truncation appeared in the designated coiled-coil (CC) domain of DksA, we investigated how the truncation could affect the fold of DksA1 using AlphaFold2 (Jumper *et al*., 2021; Skolnick *et al*., 2021). The polypeptide sequences of DksA and DksA1 were submitted to the colabfold2 server (AlphaFold2) with default settings and were modelled with a moderate-to-high degree of confidence (Figure S1C, D). The CC domain of DksA was well built as was the majority of the globular-head (GH) domain. Conversely, the eight-amino acid truncation in DksA1 resulted in a partial disruption of the tip of the CC domain with AlphaFold predicting a ‘disordered’ loop with moderate confidence (Figure S1D). The extent of the potential structural changes in the mis-formed tip of the CC domain in DksA1 can be observed when overlaid with a previously solved structure of a DksA:RNAP complex from *E. coli* (PDB ID: 5VSW) (Molodtsov *et al*., 2018). As shown in Figure 1A, all DksA variants align well, except for the altered CC-loop regions near helix ɑ3 and the C-terminal region helix ɑ5 not present in the DksA *E. coli* crystal structure. Importantly, the truncation did not delete key function residues shown to be needed for interaction of DksA with the RNAP allosteric (K104, L101, K100, R97) or with the RNAP active site (D80, D77) (Parshin *et al*., 2015; Molodtsov *et al*., 2018) (Figure 1B). Furthermore, all DksA variant structures can be modeled into RNAP (Figure S2) slotting into the allosteric site (Figure 1C) and RNAP active site (Figure 1D) without significant variation from the experimental DksA structure. However, while the AlphaFold predictions for Lp02 DksA and DksA1 had a properly modeled ppGpp binding site (Figure 1E), the twin-aspartate loop of the CC domain was significantly altered in DksA1 through the shift of D69 (D71 in DksA) by ∼8.9Å (Figure 1F). These models therefore suggest that while the functional residues are conserved from the truncation, DksA1 should be incapable of coordinating Mg^2+^ in the active site of RNAP. Conversely, the structural predictions show that DksA1 is likely still able to bind RNAP. Specifically, DksA1 is predicted to still contact the secondary binding site of ppGpp functionally as an allosteric inhibitor of RNAP through ppGpp-mediated recruitment.

**Figure 1:**
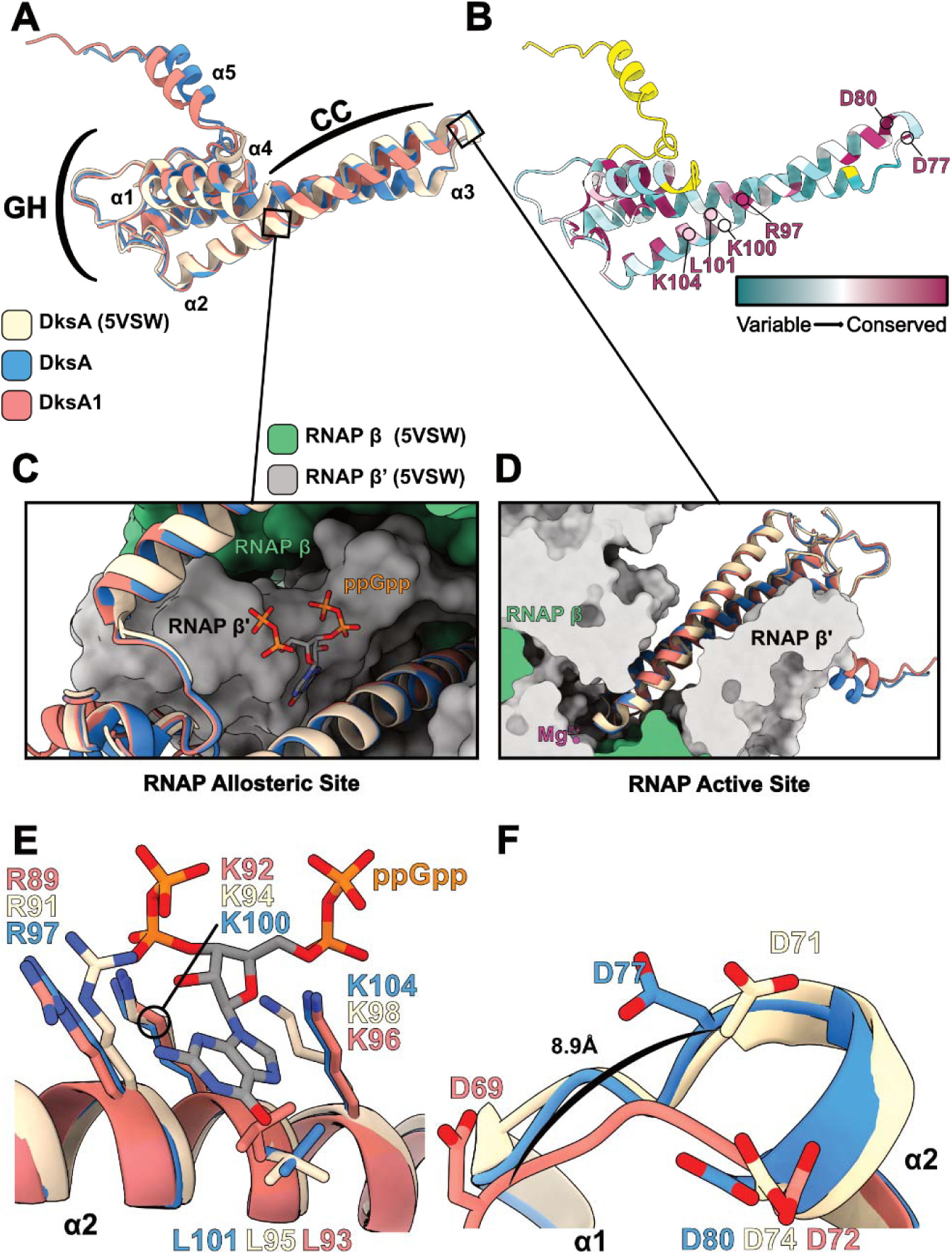
Structural comparison of DksA and DksA1 AlphaFold models. (A) Structural overlay of DksA (PDB ID: 5VSW) with both Lp02 DksA and DksA1 as modeled by AlphaFold. (B) Residue conservation of Lp02 DskA as plotted by the server Consurf (Landau *et al*., 2005; Ashkenazy *et al*., 2016). Yellow indicates insufficient information in the multisequence alignment. Indicated on the structure are conserved residues important for binding ppGpp and allostery with RNAP (R97 to K104), in addition to contact with the active site (D77-D80). (C) Binding of DksA with the RNAP allosteric site. A DksA:RNAP experimental structure (PDB ID: 5VSW) and aligned AlphaFold models are shown as cartoons, whereas RNAP is drawn as a molecular surface. The structure of ppGpp bound in the complex is also shown. (D) Slice view of DksA showing the contact point of residues D80 and D77 with magnesium in the RNAP active site. (D) Interactions of DksA variants with ppGpp in the RNAP allosteric site are shown and not disturbed by the truncation in DksA1. (E) Interactions of DksA variants with magnesium in the active site of RNAP are shown. AlphaFold predicts the truncation in DksA1 drastically moves the position of a critical aspartate residue (D69/71/77).

### PsrA partially rescues the growth defect of the *dksA1*, but not Δ*dksA*, strain in *A. castellanii*

Given the previously described regulatory roles of DksA and PsrA (Dalebroux *et al.,* 2010; Graham *et al.,* 2021), the discovery of the Δ*psrA dksA1* strain along with the *dksA1* strain presented a fortuitous opportunity to investigate potential regulatory links between the two proteins in the established *A. castellanii* protozoan infection model. Intracellular growth kinetics were conducted on *dksA1* pThy, Δ*psrA* pThy, *dksA1* Δ*psrA* pThy, and *dksA1* pVect-*psrA* strains, along with virulent Lp02 pThy and avirulent Δ*dotA* pThy strains as controls, at 25°C over 168 h as previously described (Tanner *et al.,* 2016; Graham *et al*., 2021) (Figure 2A). The profiles of Lp02 pThy and Δ*dotA* pThy strains were as expected demonstrating ∼ three-log_10_ increases in titre, and titres staying near initial inoculum concentration, respectively. The *dksA1* pThy and Δ*psrA dksA1* pThy strains showed no significant increase in titres indicating that the chromosomal single copy *dksA1* mutation was defective for virulence. Surprisingly, *in trans* PsrA expression in the *dksA1* background resulted in a two-log_10_ recovery of growth (*p* = 0.0006) to a titre within one-log_10_ of the Lp02 pThy strain. This suggested PsrA were able to partially, but not fully, rescue the avirulent phenotype demonstrated by the *dksA1* strain. While a genomic copy of *psrA* is present in the *dksA1* strain, it was shown previously that *in trans* expression of PsrA from pVect-*psrA* resulted in an elevated quantity of cellular protein that not only rescued the reduced growth phenotype associated with gene deletion of *psrA*, but also enhanced replicative growth of Lp02 in *A. castellanii* (Graham *et al*., 2021). Nevertheless, the surprising ability of PsrA to partially complement the growth defect of *dksA1* led us to expand our investigations to define the relationship between the two regulatory factors.

**Figure 2:**
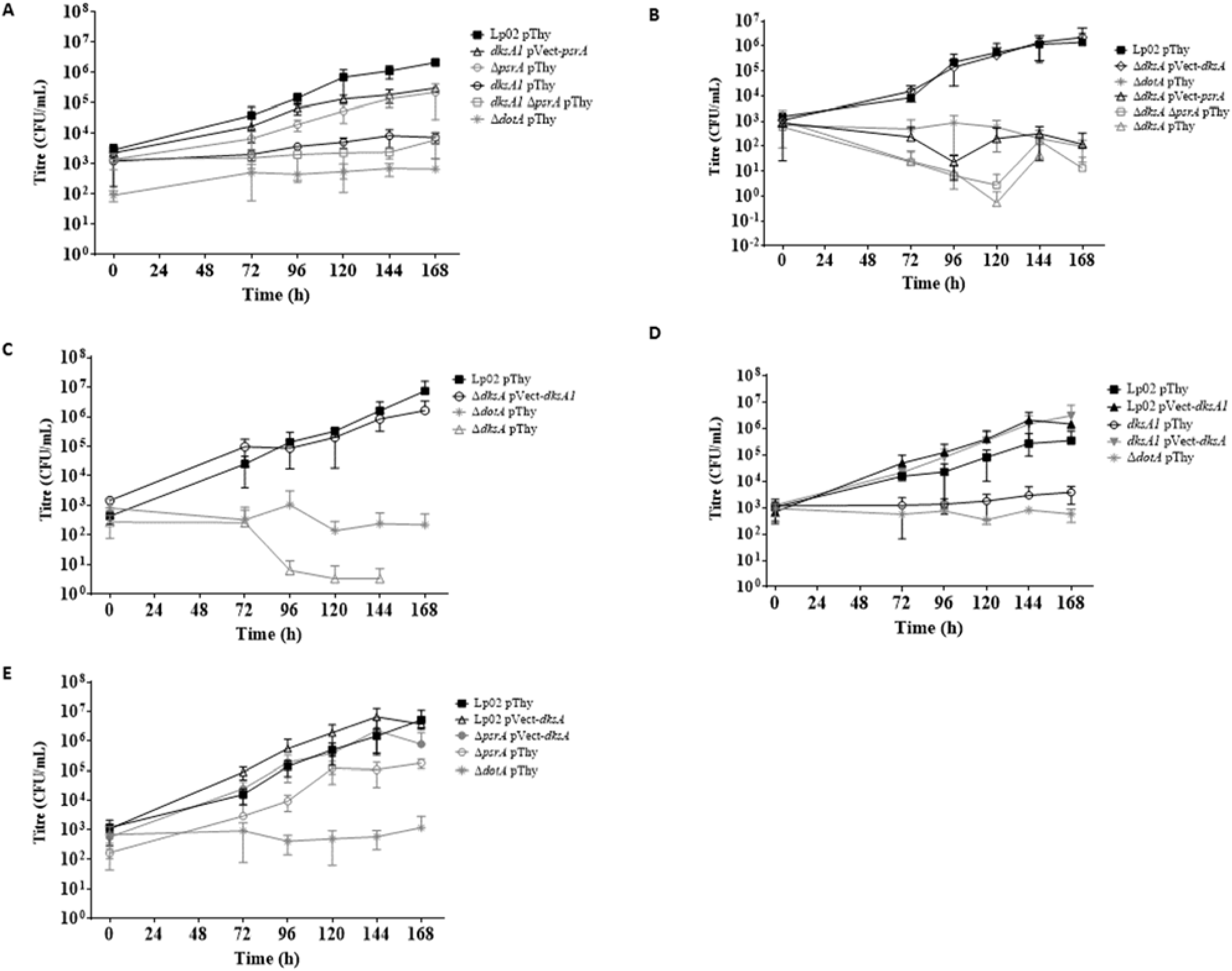
**Synergistic impacts of DksA, DksA1, and PsrA on virulence of *L. pneumophila* in the *A. castellanii* protozoa.** Intracellular growth kinetics of *L. pneumophila* strains in *A. castellanii* protozoa at 25°C. Strains tested were parental Lp02 and avirulent Δ*dotA* along with isogenic strains featuring single or combinatorial genetic mutations (Δ*psrA*, Δ*dksA*, *dksA1*) with or without *in trans* expression of *psrA, dksA* or *dksA1*. Error bars represent SD across mean of three biological replicates, with two technical replicates each. Samples below detection limit (∼10 CFU/mL) were recorded as zero and omitted from graph if they lacked countable colonies across all replicates.

A previous study elsewhere reported that an insertional *dksA* mutant showed a modest intracellular growth defect in *A. castellanii* at 37°C with slow growth initially, but eventually achieving similar titres to Lp02 by the end of the 96 h experiment (Dalebroux *et al,* 2010). This reported phenotype differs in comparison to the significantly reduced intracellular growth observed with the *dksA1* strain in this study, thereby supporting our hypothesis that DksA1 bore altered function and distinct phenotype versus the null mutant. To test this, and to determine if the compensatory effect of *psrA* observed with the *dksA1* mutant strain extended to a null deletion mutant strain, we generated single Δ*dksA* and double Δ*dksA* Δ*psrA* mutant strains. The intracellular growth kinetics of the strains were assessed in *A. castellanii* under the same conditions (Figure 2B). Interestingly, the Δ*dksA* pThy and Δ*dksA* Δ*psrA* pThy mutant strains were both fully avirulent, starting at a slightly lower titre than the Δ*dotA* pThy strain, then declining to, or below, the detection limit of the assay (∼10 CFU/mL). The severe growth defect of the Δ*dksA* pThy strain was fully rescued by *in trans* expression of *dksA*, but not *psrA*; a notable divergence from the rescued phenotype demonstrated by the *dksA1* pVect-*psrA* strain. Taken together, the lack of DksA results in a complete avirulent phenotype in these conditions which cannot be rescued by PsrA, whereas the hypomorph DskA1 has a less severe impact on intracellular growth that can be partially rescued by PsrA.

### DksA1 rescues the growth defect of the Δ*dksA* strain, and vice versa, in *A. castellanii*

Since DksA can rescue the severe growth defect of the Δ*dksA* strain in *A. castellanii* (Figure. 2B), we next asked if the partially functional DksA1 could likewise do so despite the truncation in the coiled-coil structure (Figure 1). To answer this, the Δ*dksA* pVect-*dksA1* strain was assessed in *A. castellanii* along with Δ*dksA* pThy, Lp02 pThy and Δ*dotA* pThy (Figure 2C). The severely reduced growth phenotype of the Δ*dksA* pThy strain was robustly rescued by *in trans* expression of DksA1 as shown by the growth profile of the Δ*dksA* pVect*-dksA1* strain (*p* = 0.0002) matching that of the Lp02 pThy strain. It should be noted that this growth phenotype contrasted with the minimal growth profile of the *dksA1* pThy strain (Figure 2A), suggesting that restoration of the virulent phenotype of the Δ*dksA* strain was due to *in trans* expression of DksA1 protein (pVect*-dksA1*) of sufficient levels to offset the structural and functional inefficiency. Not unexpectantly, *in trans* expression of DksA successfully rescued the avirulent phenotype of the *dksA1* pThy strain to match the growth profile of Lp02 pThy most likely due to the *in trans* expression of DksA of which levels exceeded that of DksA1 (Figure 2D). In the reverse scenario, the growth profile of Lp02 pVect-*dksA1* still matched that of Lp02 pThy suggesting that despite the *in trans* expression of DksA1, the chromosomal-copy production levels of DksA and PsrA in parallel were each sufficient to disguise any effects caused by the dysfunctional DksA1 (Figure 2A, Figure 2D).

### DksA restores the growth defect of the Δ*psrA* strain in *A. castellanii*

To further define the relationship between DksA and PsrA, we next asked if DksA could reciprocally compensate for lack of PsrA or generally, affect intracellular growth in the presence of PsrA. To answer this, Lp02 pVect-*dksA* and Δ*psrA* pVect-*dksA* strains were assessed for growth in *A. castellanii* along with the Lp02 pThy, Δ*dotA* pThy and the Δ*psrA* pThy strains for comparative purposes (Figure 2E). As previously reported, the Δ*psrA* pThy strain consistently maintained 1 log_10_ lower than that of Lp02 pThy (*p =* 0.029) (Graham *et al*., 2021). The mean growth profile of the Lp02 pVect-*dksA* strain showed a subtle increase of ∼½ log_10_ over that of Lp02 pThy (*p* = 0.007); this enhancement could also be seen in the <u>Δ</u>*dksA* pVect*-dksA* complemented strain (Figure 2D; *p* = 0.0247). Likewise, we found that *in trans* expression of DksA in the Δ*psrA* pVect-*dksA* strain rescued the growth defect of Δ*psrA* strain, increasing growth by ∼1 log_10_ to match that of Lp02 pThy strain (p < 0.001 versus Δ*psrA* pThy, not significant versus Lp02 pThy). Although this experiment did not generate large phenotypic differences, we found the tendency towards subtle phenotypes upon *in trans* expression of DksA to be noteworthy relative to the very strong phenotypes noted in the *dksA* mutant strains.

### DksA is not required for intracellular growth in human U937-derived macrophages

Some *L. pneumophila* genes exhibit host specific requirements such that a subset are required only for growth in one host type, but not in another. Previous work showed that the human U937-derived macrophage infection model is more permissive to *L. pneumophila* regulatory mutant strains than *A. castellanii*, as exemplified by transcriptional regulator Δ*psrA* or Δ*cpxRA* TCS mutants (Tanner *et al.,* 2016; Graham *et al.,* 2021). DksA was previously shown to be dispensable for growth in primary murine bone marrow macrophages (Dalebroux *et al.,* 2010).

However, since the phenotype we observed for the Δ*dksA* strain in *A. castellanii* was different from that reported by Dalebroux *et al*. (2010), we decided to ascertain the phenotype using the human monocytic U937 cell-line as a representative mammalian cell model. Intracellular growth kinetics were assessed for the same set of strains previously used in the *A. castellanii* infection assays (Figure 2). As expected, Lp02 pThy grew well, increasing approximately 100-fold by the 72 h timepoint (Figure 3A), and Δ*dotA* pThy maintained a steady titre with no growth. In comparison to that of Lp02 pThy, Δ*dksA* pThy and Δ*dksA* Δ*psrA* pThy showed similarly robust, but offset growth profiles. Additionally, despite normalization of the inoculum by OD_600_, these two strains showed initial quantified titres lower than that of Lp02 pThy (Figure 3A). Only Δ*dksA* Δ*psrA* pThy showed a significant deviation from Lp02 pThy (*p* = 0.049). The Δ*dksA* pVect-*psrA* showed a similar profile to that of Lp02 pThy (Figure 3A), and did not show the reduced starting titre, but the difference to Δ*dksA* pThy was not significant. No significant change in phenotype was observed with the remaining set of strains featuring *in trans* DksA expression, either by complementation (Δ*dksA* pVect-*dksA*), overexpression, (Lp02 pVect-*dksA*) or in Δ*psrA* pVect-*dksA* (Figure 3B). The growth profiles of the *dksA1* pThy, *dks1* Δ*psrA* pThy, or *dksA1* pVect-*psrA* were indistinguishable from Lp02 pThy (Figure 3C) and lacked even variability in the initial uptake. Thus, these results verify that DksA is not required for, nor does it enhance, intracellular growth in U937-derived macrophages.

**Figure 3:**
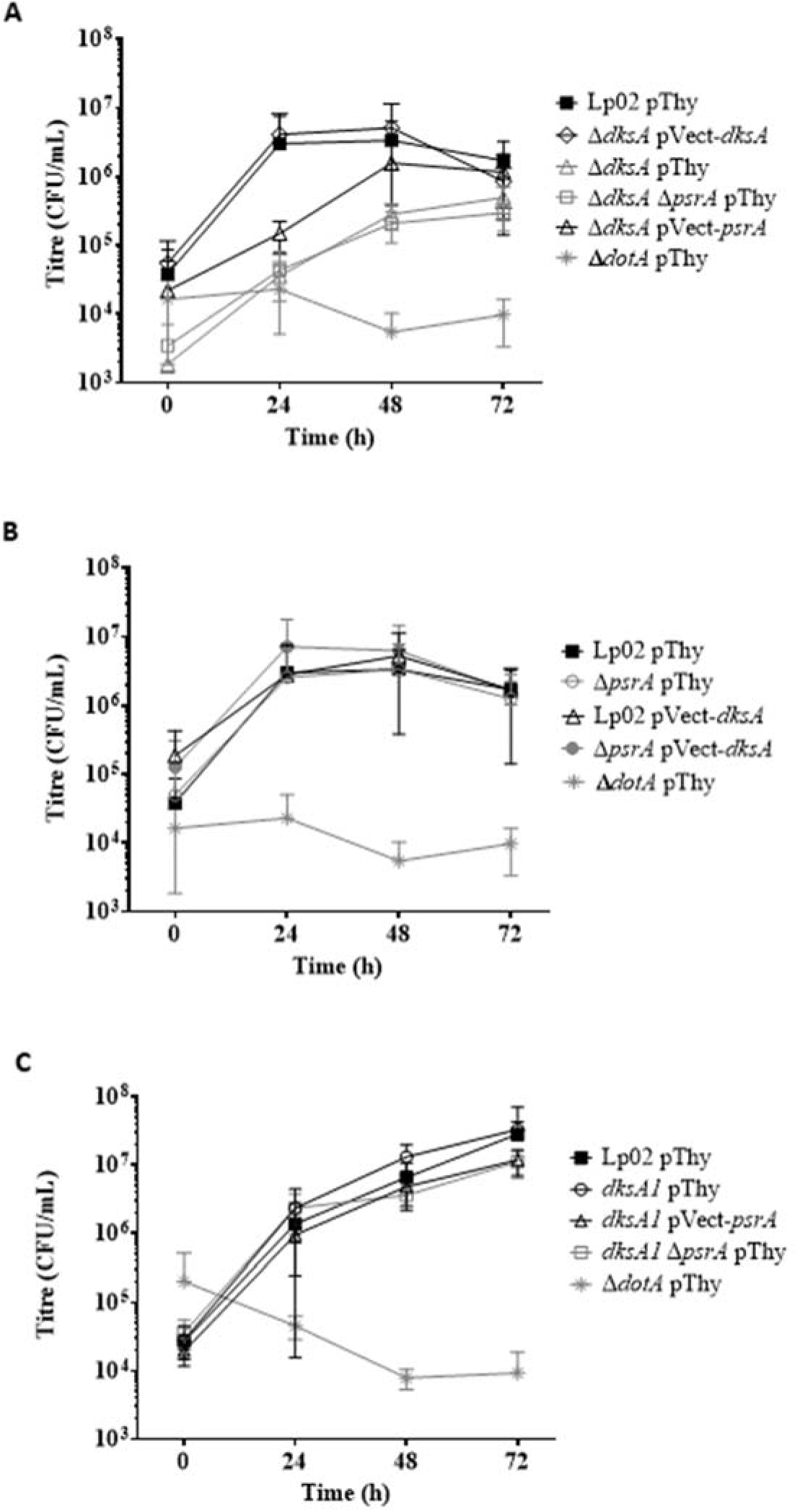
DksA is dispensable for growth in human macrophages. Intracellular growth kinetics of *L. pneumophila* strains in U937-derived macrophages at 37°C. Strains tested were parental Lp02 and avirulent Δ*dotA* along with isogenic strains featuring single or combinatorial genetic mutations (Δ*psrA*, Δ*dksA*, *dksA1*) with or without *in trans* expression of *psrA, dksA* or *dksA1*. Error bars represent SD across mean of three biological experiments with two technical replicates each. Experiments in panels A-C were conducted concurrently, but presented separately for clarity.

### *In vitro* growth is impacted by DksA, PsrA and DksA1

The differences observed with the intracellular growth kinetics of isogenic mutant strains with and without *in trans* gene expression constructs in *A. castellanii* may be in part due to defects in the growth rates of the cells, rather than solely virulence defects *in vivo*. To distinguish this possibility and more generally characterize the *in vitro* growth characteristics of the strains, we tested the *in vitro* growth rates of the strains in BYE broth at 37°C over a 24-hour time period (Figure 4). While the growth profiles of *dksA1* pThy and Lp02 pThy were similar to one another, both were distinct from the growth profile of Δ*psrA* pThy which was significantly reduced in comparison to that of *dksA1* pThy (*p* = 0.0092) and Lp02 pThy (*p* = 0.0007) (Figure 4A) with the latter comparison consistent with our previous study (Graham *et al*., 2021). The growth profiles of Lp02 pThy, *dksA1* pVect- *psrA*, and *dksA1* Δ*psrA* pThy were all similar to one another indicating that lack of PsrA, or elevated levels of PsrA from *in trans* expression, did not affect the growth profile of the *dksA1* (Figure 4B). Interestingly, the *in vitro* growth kinetics of Δ*dksA*-based strains featured growth variability across replicates as manifested by the large error bars for hourly data points (Figures 4C and 4D). Despite this variability, the growth profile of Δ*dksA* pThy was not considered to be significantly different from that of Lp02 pThy (Figure 4C). Notably, the growth variability disappeared from the Δ*dksA* pVect-*dksA* growth profile suggesting this was a complementable phenomenon (Figure 4C). *In trans* expression of *dksA1* (Δ*dksA* pVect-*dksA1*) significantly reduced the growth profile in comparison to that of Lp02 pThy (*p* = 0.0053) indicating the elevated levels of DksA1 compromised replicative growth (Figure 4C). As observed with *dksA1* Δ*psrA* pThy (Figure 4B), the growth profile of Δ*dksA* Δ*psrA* pThy was similar to that of Lp02 pThy (Figure 4D) indicating the dispensability of PsrA. *In trans* expression of DksA in Lp02 (Lp02 pVect-*dksA*) did not alter the growth profile from that of Lp02 pThy (Figure 4E). While a significant reduction was observed for the growth profile of Δ*psrA* pVect-*dksA* in comparison to that of Lp02 pThy (*p* = 0.0053; Figure 4E), the difference was not due to *in trans* expression of DksA as a similar difference was observed for Δ*psrA* pThy versus Lp02 pThy (Figure 4A). *In trans* expression of PsrA did not reduce the growth variability phenotype of Δ*dksA* nor was the growth profile significantly different from that of Lp02 pThy (Figure 4F). *In trans* expression of DksA1 had no effect as the growth profiles of Lp02 pVect-*dksA1* and Lp02 pThy were similar (Figure 4E).

**Figure 4:**
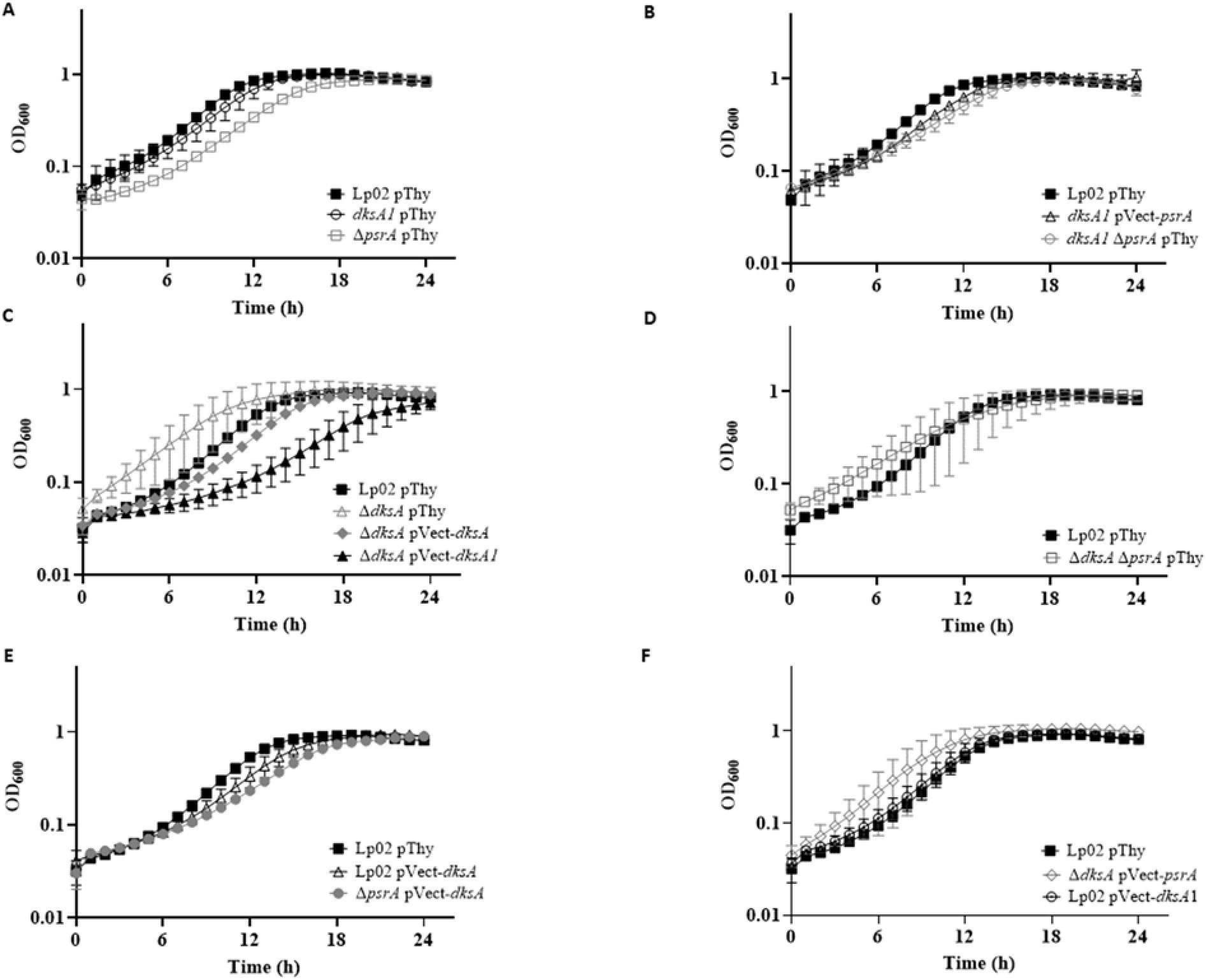
***L. pneumophila in vitro* growth at 37°C is impacted by DksA, PsrA and DksA1.** *In vitro* growth kinetics of parental Lp02 and isogenic strains featuring single or combinatorial genetic mutations (Δ*psrA*, Δ*dksA*, *dksA1*) with and without *in trans* expression of *psrA, dksA* or *dksA1* genes. Plate-grown bacteria were suspended in BYE broth and normalized to initial OD_600_ = 0.15, and growth at 37°C was monitored for OD_600_ hourly for 24 h. Graphs in panels A-F were experiments conducted concurrently, but presented separately for clarity. Error bars are SD across mean of three independent biological replicates with three technical replicates each.

The *in vitro* growth profile assays were repeated at 25°C, albeit over a 48-hour time period to account for the slower growth rate, to mimic the incubation temperature of the *A. castellanii* assays. The resultant growth profiles were mostly similar to those attained at 37°C, with growth variability of Δ*dksA* pThy exacerbated at 25°C as indicated by data points with widened error bars (Figure 5C). One notable difference was that the growth profile of Δ*dksA* pVect-*dksA1* matched that of Lp02 pThy at 25°C (Figure 5D), versus the reduction at 37°C (Figure 4C), suggesting that the slower growth rate and/or lower temperature accommodated the dysfunctional nature of DskA1.

**Figure 5:**
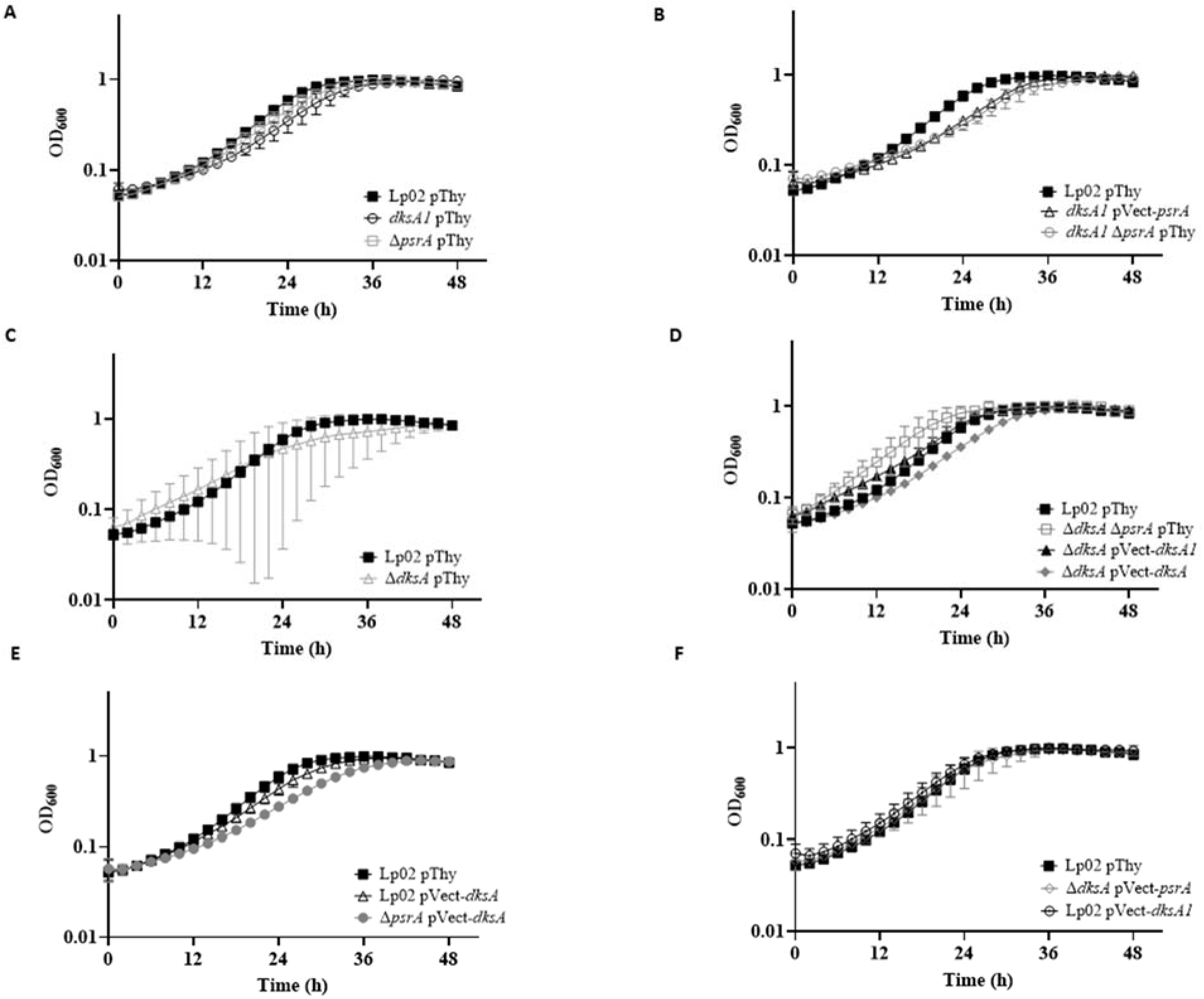
***L. pneumophila in vitro* growth at 25°C is impacted by DksA, PsrA and DksA1.** *In vitro* growth kinetics of parental Lp02 and isogenic strains featuring single or combinatorial genetic mutations (Δ*psrA*, Δ*dksA*, *dksA1*) with and without *in trans* expression of *psrA, dksA* or *dksA1* genes. Plate-grown bacteria were re-suspended in BYE broth and normalized to initial OD_600_ = 0.15, and growth at 25°C was monitored for OD_600_ every 2 h for 48 h. Graphs in panels A-F were experiments conducted concurrently, but presented separately for clarity. Error bars are SD across mean of three independent biological replicates with three technical replicates each.

These findings together suggest that while strains lacking *dksA* featured more variability in growth profiles, more so at 25°C then at 37°C, the *in vitro* growth assays (Figures 4 and 5) illustrates that the observed *in vivo* growth defects are specific to *A. castellanii* (Figure 2), and not due to poor growth of the bacterial cells themselves.

### DksA, and possibly PsrA, affects cell morphology

Throughout this study, it was noted that the colony morphologies of Δ*dksA* and *dksA1* strains grown on BCYE agar plates differed significantly from that of Lp02. In comparison to the opaque beige-white glossy colonies with varied margins featured by Lp02, Δ*dksA* colonies were textured matte white with defined margins, and *dksA1* colonies were smooth glossy white with defined margins. The phenotypes were particularly evident when bacterial suspensions in broth were spotted onto BCYE agar plates as were done in serial dilution assays in this study (Figure 6). The phenotype was fully rescued by *in trans* expression of DksA for both Δ*dksA* and *dksA1* strains, whereas *in trans* DksA1 expression only partially rescued the phenotype of the Δ*dksA* strain. The colony morphology of the Δ*dksA* Δ*psrA* strain was similar to that of the Δ*dksA* strain, but not to the Δ*psrA* strain which was similar to that of Lp02 indicating the sole influence of DksA. Finally, *in trans* PsrA expression did not appear to alter colony morphology in all strain backgrounds assessed (Lp02, Δ*dksA*, *dksA1,* Δ*psrA*) confirming that PsrA does not influence this phenotype.

**Figure 6:**
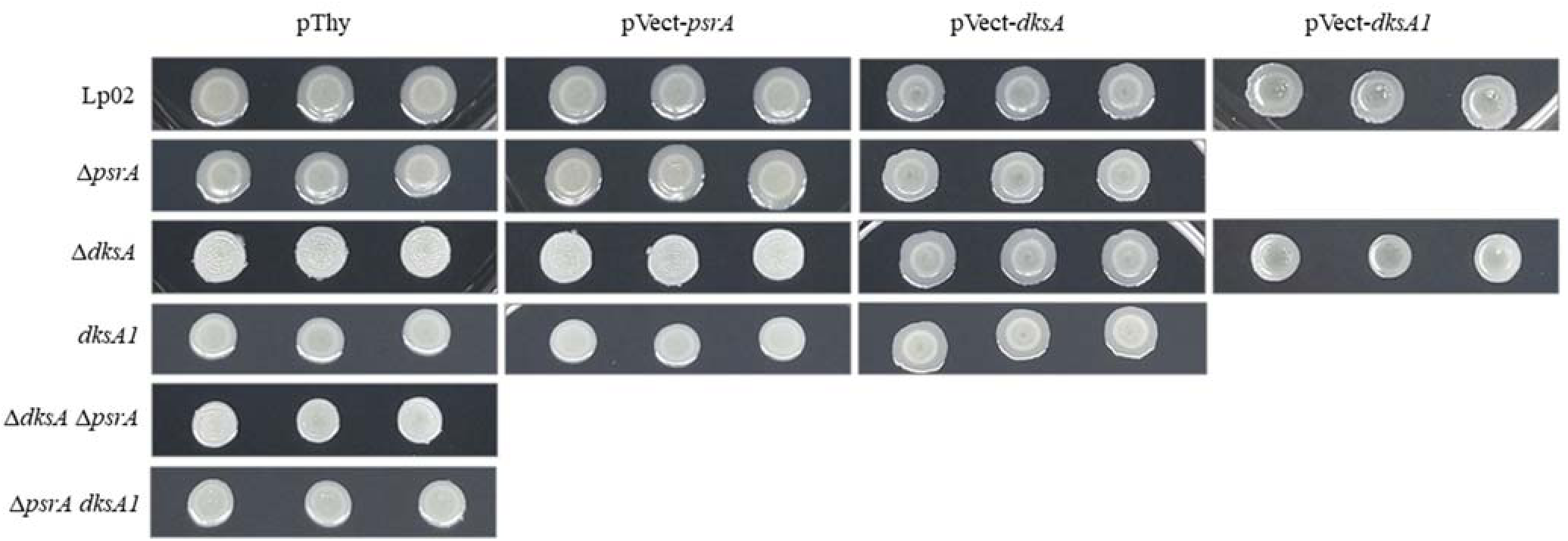
***L. pneumophila* DksA and DksA1, but not PsrA affects colony morphology.** Parental Lp02 and isogenic strains featuring single or combinatorial genetic mutations (Δ*psrA*, Δ*dksA*, *dksA1*) with and without *in trans* expression of *psrA, dksA* or *dksA1* genes. Strains were grown on BCYE plates for 3 days and used to generate a bacterial suspension in BYE at OD_600_ = 0.15. 10 μL of the suspension was spotted onto BCYE plates in triplicate then the plates were incubated at 37°C at 5% CO_2_ for 4 days prior to being photographed. Images are representative of one of three biological replicates.

We previously reported that PsrA promoted cell elongation when overexpressed *in trans* (Graham *et al.,* 2021). Conversely, it was reported elsewhere that lack of DksA influenced cell elongation (Dalebroux *et al*., 2010). To assess whether a genetic relationship between PsrA and DksA was evident in cell structure morphology, and to further explore whether *dksA1* generated a phenotype distinct from Δ*dksA*, bacteria in exponential growth phase (EP) and in stationary phase (SP) were microscopically imaged (Figure 7) and analyzed for cell lengths (Figures 8A and 8B). As a control, Lp02 pThy bacteria formed typical rods (mean length 3.6 µm) in EP, that elongated into SP (mean length 5.8 µm). The Δ*dksA* pThy bacteria were notably elongated in EP (mean length 6.2 µm) and, in SP, became extremely elongated into filamentous forms often of sufficient length to often exceed the field of view, averaging 13.2 µm among measurable cells (with the entire cell visible in one field of view). Likewise, the Δ*dksA* Δ*psrA* pThy bacteria were also elongated in EP, and strongly filamentous in SP (mean lengths 5.4 µm and 17.1 µm, respectively). Conversely, Δ*psrA* pThy bacteria were morphologically indistinguishable from that of Lp02 pThy as previously observed (Graham *et al*, 2021). The filamentation of Δ*dksA* pVect-*dksA* bacterial forms was rescued by *in trans* DksA expression as they were slightly, but significantly shortened relative to Lp02 (2.9 µm and 4.7 µm in EP and SP phases, respectively). Expression of DksA *in trans* in Δ*psrA* pVect-*dksA* and Lp02 pVect-*dksA* strains also resulted in slightly, but significantly shorter cell lengths in comparison to those of Lp02 pThy and comparable to Δ*dksA* pVect-*dksA* in both growth phases. *In trans* PsrA expression in Δ*dksA* pVect-*psrA* did not strongly affect the bacterial cell lengths and elongation was still present (7.5 µm and 14.2 µm in EP and SP, respectively). These findings suggest that the shortening is mediated by DksA independently of PsrA.

**Figure 7:**
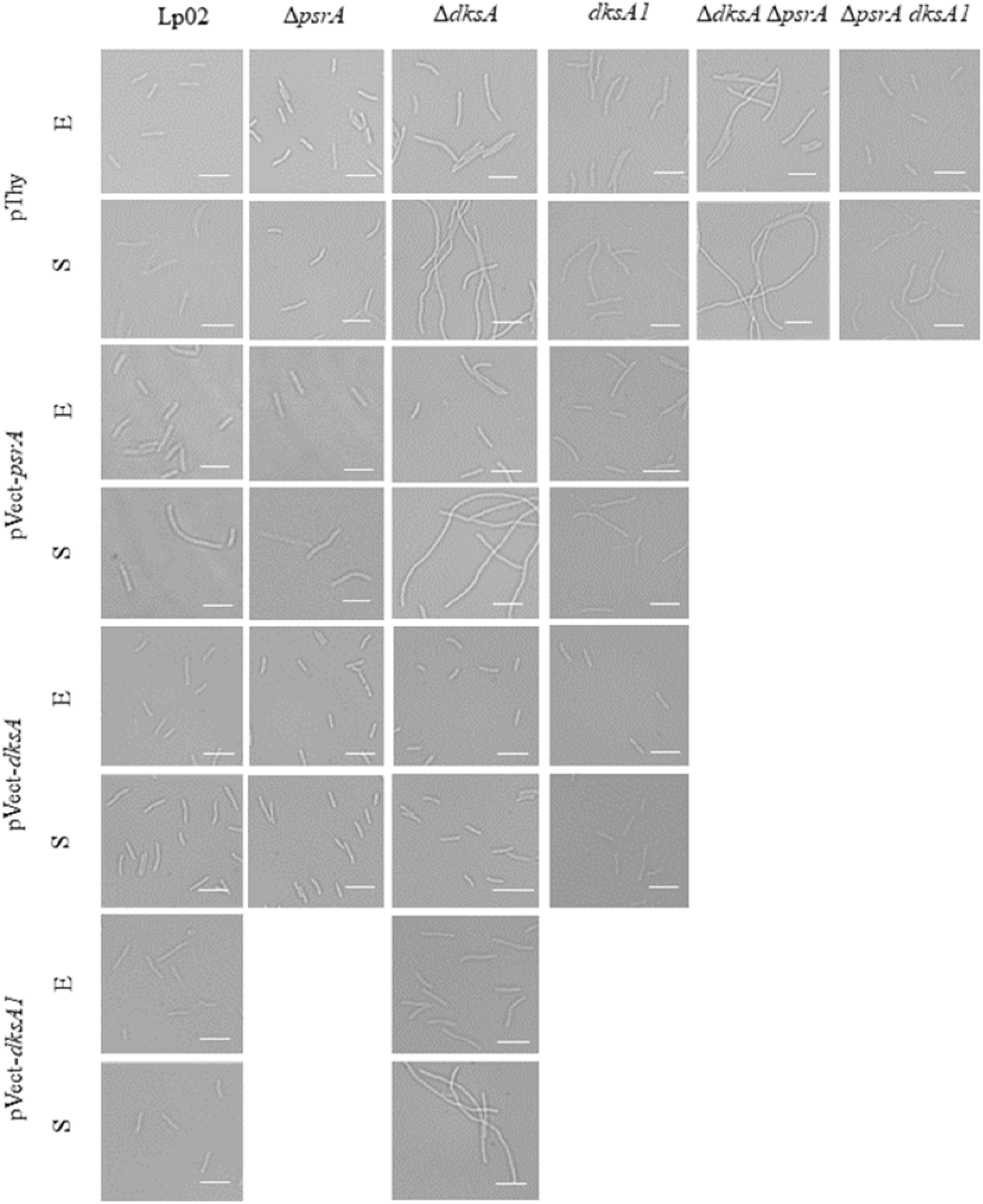
***L. pneumophila* Δ*dksA*, but not *dksA1* or *psrA* cause filamentation.** Parental Lp02 and isogenic strains featuring single or combinatorial genetic mutations (Δ*psrA*, Δ*dksA*, *dksA1*) with or without *in trans* expression of *psrA, dksA* or *dksA1* genes were grown on BCYE plates and used to inoculate a dilute suspension of OD_600_ = 0.02 in BYE broth then incubated overnight at 37°C with aeration, to exponential (E) (∼18h , OD_600_ = ∼0.5) or stationary (S; 24 h after EP) growth phases, then live-mounted on a 2% agarose pad prepared in ddH_2_O and imaged on an Axio Observer Z1 inverted microscope (Zeiss) equipped with a glycerol-immersion 150X objective. Representative microscopic images of bacteria from one of three biological replicates. Scale bar is 5 µm.

**Figure 8:**
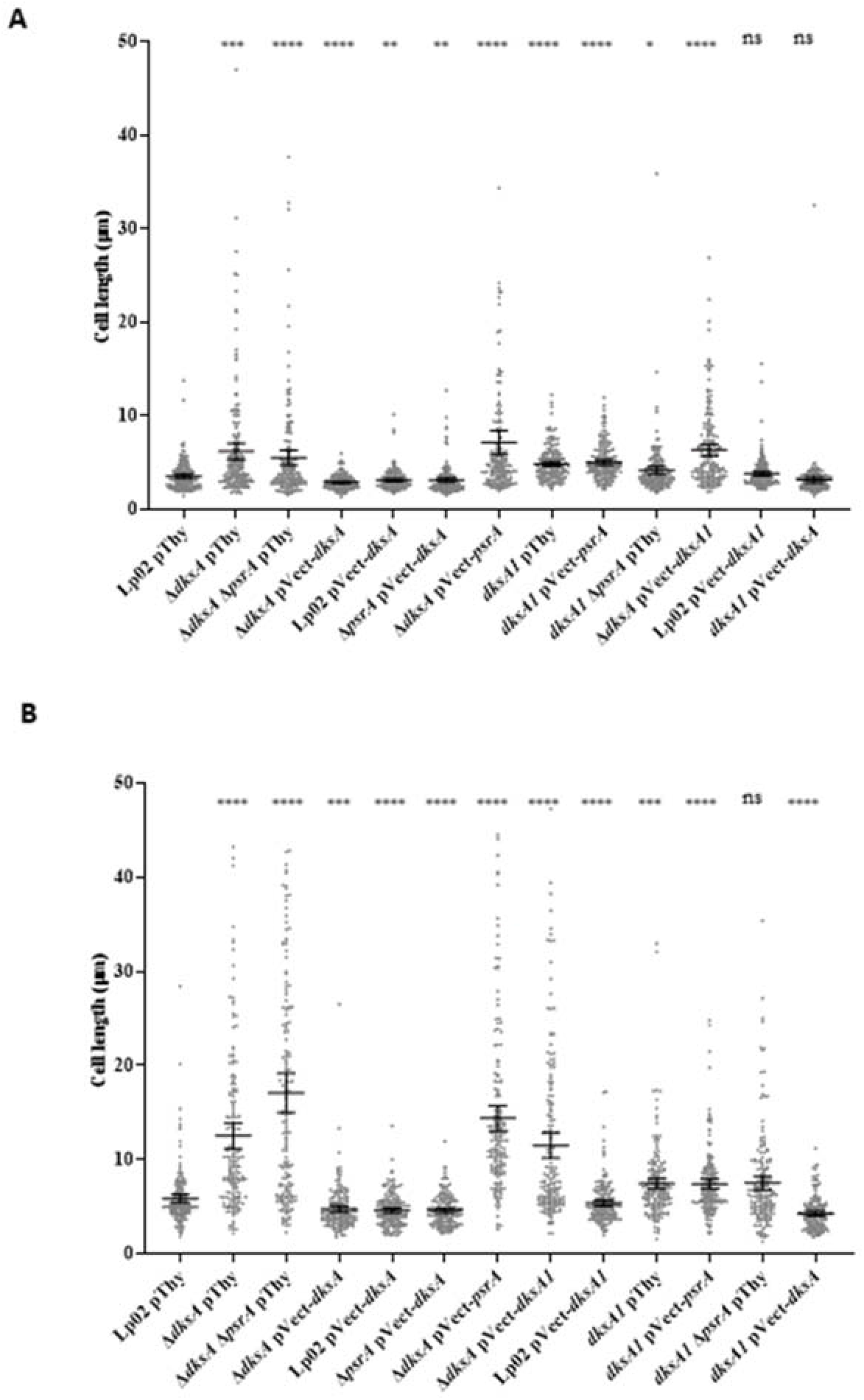
***L. pneumophila* Δ*dksA*, *dksA1*, and *psrA* contribute to cell length regulation.** Parental Lp02 and isogenic strains featuring single or combinatorial genetic mutations (Δ*psrA*, Δ*dksA*, *dksA1*) with or without *in trans* expression of *psrA, dksA* or *dksA1* genes were grown from 3-day old plates subcultured to BYE broth at OD_600_ = 0.02 then incubated at 37°C with aeration to (A) exponential (overnight, OD_600_ ∼ 0.5) or (B) stationary phases (24 h after exponential phase) then imaged with an Axio Observer Z1 inverted microscope (Zeiss) equipped with a glycerol-immersion 150X objective and individual bacterial cell lengths quantified using the length tool in ImageJ. Where filamentation was evident, only individual cells for which the full length was visible were measured. Data represents >150 individual readings spread equally across three biological replicates. Bar on graph and error bars represents mean and 95% confidence interval, respectively. Statistical significance determined by Student’s T-test (*, *p* < 0.05; ** *p* < 0.01; *** *p* < 0.001; **** *p* < 0,0001; ns, not significant, *p* > 0.05).

The bacterial forms of *dksA1* pThy, *dksA1 ΔpsrA* pThy, and *dksA1* pVect-*psrA* strain were all slightly elongated relatively to Lp02 pThy in both EP (5.0 µm, 5.1 µm and 4.3 µm, respectively) and SP (7.5 µm, 7.7 µm, and 7.8 µm, respectively), but were not considered filamentous, and did not vary significantly from each other supporting the hypothesis that the *dksA1* allele maintained partial function. Interestingly, the Δ*dksA* pVect-*dksA1* strain did not show rescue of the elongation to resemble *dksA1* levels as might be expected (5.6 µm in EP, 12.3 µm in SP). To rule out the simple possibility of the two gene products conflicting in some way, we tested the *dksA1* pVect-*dksA* (plasmid- and chromosomal based alleles reversed). The bacterial forms of this strain appeared somewhat shortened relative to Lp02 pThy in SP only (4.2 µm) resembling the other *in trans* DksA expression strains, suggesting that relative protein concentrations may drive the phenotype. Together, this indicates that the previously reported role of PsrA is overshadowed by the more potent role of DksA, and that the DksA1 protein shows an intermediate phenotype illustrating that DksA1 is a partially functional protein in these strains.

### DksA, or DksA1, is necessary for extended culturability in depleted media

Previously, the insertional mutant Δ*dksA* strain was also noted to feature reduced recovery of culturable bacteria after extended incubation in depleted media, which the authors referred to as “survival” (i.e. culturability) (Dalebroux *et al.,* 2010). Over the course of this study, we noticed that the Δ*dksA* mutants could not be consistently sub-cultured, or often failed to grow entirely, if taken from BCYE agar plates grown more than three days. In view of this, we asked if this was a consequence of previously observed culturability phenotype occurring in the Δ*dksA* strain, and potentially the *dksA1* strains, and whether PsrA may also contribute. As this phenotype had not been assessed in the prior PsrA study (Graham *et al*., 2021), we decided to include Δ*psrA* pThy, Δ*psrA* pVect-*psrA* and Lp02 pVect-*psrA* strains from Graham *et al*. (2021).

Over the time course, optical density values peaked comparably for all strains at 24 h, and held steadily to the 96 h timepoint (Figure 9). Enumeration of titres for Lp02 pThy and Δ*psrA* pThy were comparable to one another (Figure 9A). The titres for Δ*dksA* pThy (Figure 9C) and Δ*dksA* Δ*psrA* pThy (Figure 9A) were significantly reduced by 72 h, and more so by the 96 h for Δ*dksA* pThy (Figure 9C), noting that titre enumeration was not possible for Δ*dksA* Δ*psrA* pThy at 96 h as no colonies were recovered in any replicate for this strain at that timepoint (detection limit ∼10^2^ CFU/mL; Figure 9A). The reduced culturability phenotype was rescued by *in trans* expression of DksA (Figure 9B) or DksA1 (Figure 9C), but not PsrA (Figure 9B). The extended culturability of *dksA1* pThy strain was unaffected in comparison to Lp02 pThy, thereby not requiring nor significantly benefiting from *in trans* expression of DksA or PsrA (Figure 9D). The unaltered phenotype of the *dksA1* Δ*psrA* pThy in comparison to the *dksA1* pThy strain further supports the observation that PsrA does not contribute to extended culturability (Figure 9E). Further, *in trans* expression of DksA in the Lp02 strain does not significantly improve extended culturability (Figure 9F). Thus, in summary, PsrA plays no role in extended culturability, while DksA is essential for the culturability of Lp02 in depleted media as reported elsewhere previously (Dalebroux *et al.,* 2010). Notably, the hypomorph *dskA1* allele has retained this essential function of promoting extended culturability.

**Figure 9:**
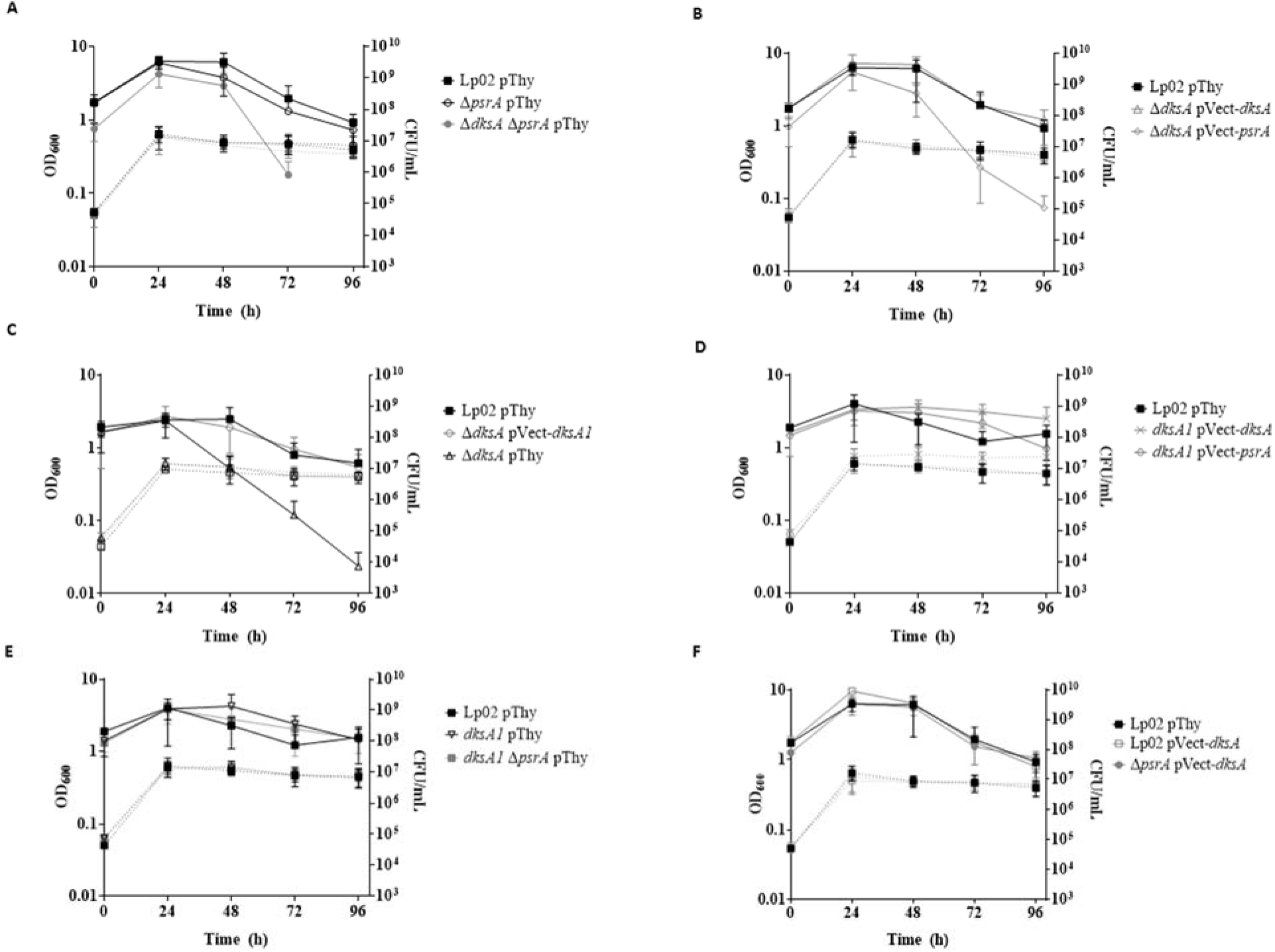
Bacterial culturability in depleted nutrient media is dependent on DksA or DksA1, but not PsrA. Panels (A-F) show *in vitro* growth kinetics of Lp02 and isogenic strains featuring single or combinatorial genetic mutations (Δ*psrA*, Δ*dksA*, *dksA1*) with or without *in trans* expression of *psrA, dksA* or *dksA1* genes in BYE broth at 37°C for 96 hours with OD_600_ values (dotted lines; left Y axis) and titre enumerated by serial dilution and incubation on BCYE plates (solid lines; right Y axis) at 24 h intervals. Data are presented as mean and SD of three biological replicates. Points were omitted if results could not be calculated (no colonies recovered, below detection limit of ∼100 CFU/mL). Graphs in panels A-F were experiments conducted concurrently, but presented separately for clarity.

### Pigmentation requires fully functional DksA, but not PsrA

*L. pneumophila* pigment is a brown, melanin-like secreted protein that is released into the supernatant during the transition to stationary phase under the control of the stringent response through RelA (Zusman *et al*., 2002; Dalebroux *et al*., 2010). Further, we observed hyperpigmentation of a subset of parental, isogenic (Δ*psrA*, Δ*dksA*) and *dksA1* strains producing DksA *in trans* (Figure 10A). To ascertain whether DksA1 and/or PsrA could contribute to this hyperpigmentation, we quantified the pigmentation for each strain as done elsewhere (Wiater *et al*., 1994; Dalebroux *et al.,* 2010) (Figure 10B).

**Figure 10:**
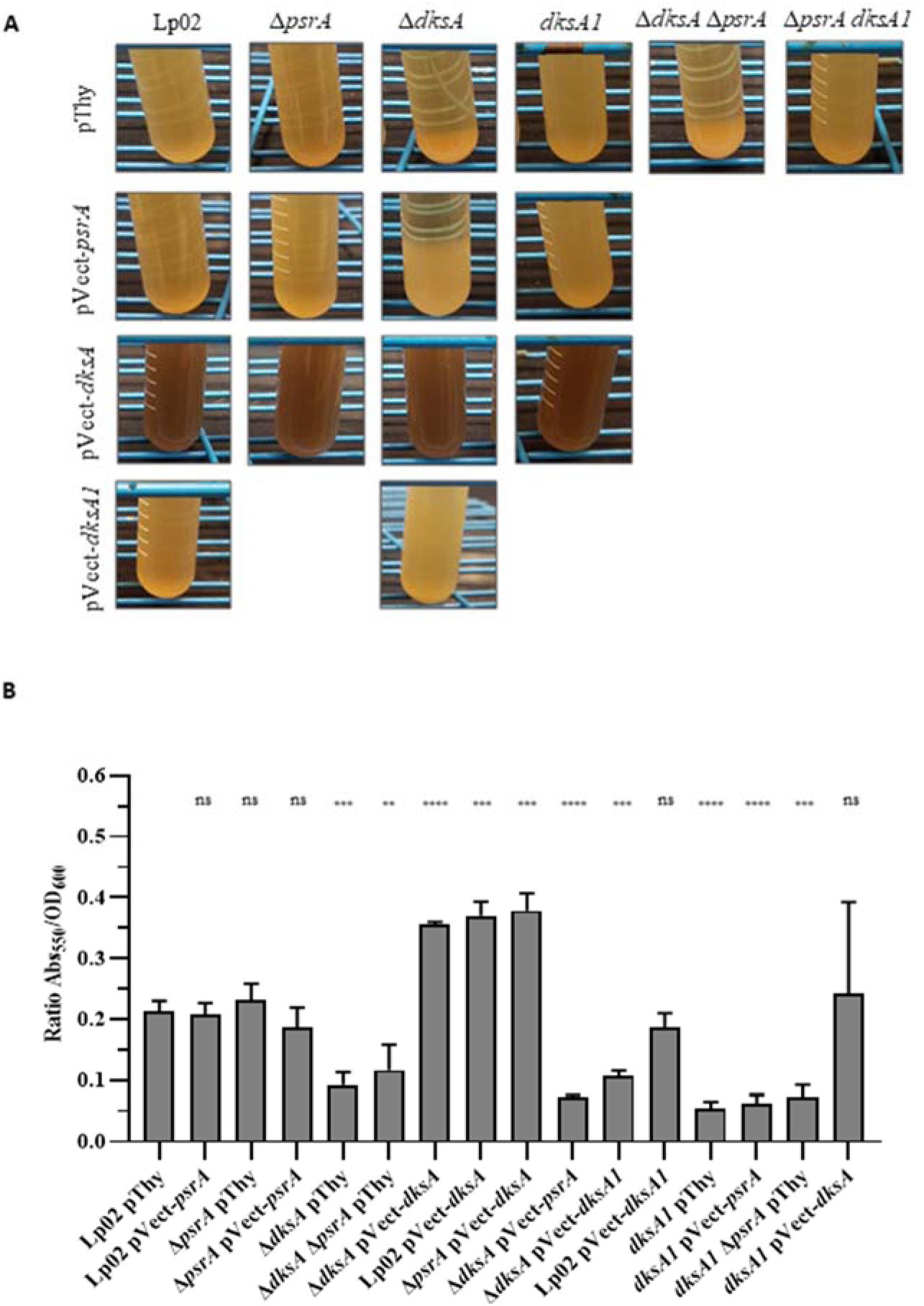
Pigmentation is regulated by DksA, but not PsrA. Parental Lp02 and isogenic strains featuring single or combinatorial genetic mutations (Δ*psrA*, Δ*dksA*, *dksA1*) with or without *in trans* expression of *psrA, dksA* or *dksA1* genes, were grown on BCYE plates for 3 days and used to generate a 5 mL bacterial suspension (BYE at OD_600_ = 0.15) then incubated at 37°C for 30 h (a) Qualitative observation of pigmentation. After 48 h incubation tubes were set aside for 24 h at room temperature to allow cells to sediment out, then photographed. Representative image of three replicates. (b) Quantification of pigment at 30 h. After 30 h of incubation 1 mL of culture was centrifuged at 20,000 x *g*. The supernatant was measured for pigment levels via absorbance at 550 nm, then the pellet was resuspended in same volume of 1X PBS pH 7.0 to measure cell density via OD_600_. Readings were taken with a plate reader in a 96 well plate relative to appropriate BYE or PBS blanks. Readings reflect Abs_550_ /OD_600_. Error bars represent SD across three biological replicates. Statistical significance calculated via Student’s T-Test and relative to parental Lp02 pThy, indicated above bars (*, *p* < 0.05, ** *p* < 0.01; *** *p* < 0.001; **** *p* < 0,0001; ns, not significant, *p* > 0.05).

Lp02 pThy strain developed moderate pigmentation (Abs_550_/OD_600_ = ∼0.20) that was similarly achieved by Δ*psrA* pThy, Δ*psrA* pVect-*psrA*, and Lp02 pVect-*psrA* strains suggesting that PsrA does not influence pigmentation (Figure 10B). This supposition is further borne out by the significantly reduced pigmentation (Abs_550_/OD_600_ = 0.10 - 0.12) featured by both of the Δ*dksA* Δ*psrA* pThy and Δ*dksA* pThy strains. Significantly increased levels (i.e. hyperpigmentation) was achieved by the Δ*dksA* pVect-*dksA* strain (Abs_550_/OD_600_ = 0.36 - 0.38) which exceeded that of Lp02 pThy likely due to *in trans* production of DksA (Figure 10B).

Indeed, the pigmentation phenotype was rescued by *in trans* DksA expression in the Δ*psrA* strain, but not by *in trans* PsrA expression in the Δ*dksA* strain, clearly indicating that only DksA positively influences pigmentation (Figure 10B). Pigmentation was also significantly and equitably reduced (Abs_550_/OD_600_ = 0.05 - 0.09) for the *dksA1* pThy and *dksA1* Δ*psrA* strains. As the phenotype was not rescued by *in trans* PsrA expression in the *dksA1* pThy strain, this further supports the lack of involvement in pigmentation by PsrA (Figure 10B). Surprisingly, the hypomorph DksA1 could not rescue pigmentation in the Δ*dksA* strain, but pigmentation of *dksA1* strain could be induced by *in trans* DksA expression. The *dksA1* pVect-*dksA* strain showed delayed onset of hyperpigmentation such that only one of three replicates had fully pigmented by 30 h, with all replicates of this strain pigmented by 48 h (Figure 10A), but at 30 h the difference was not statistically significant. These findings indicate that the ability of DksA1 to interact with RNAP is impaired as suggested by structural modelling (Figures 1, 10B). Taken together, these results indicate that DksA is sufficient to drive pigmentation independently of PsrA.

### Growth in long chain fatty acids is mediated by PsrA and DksA

The activation of the biphasic switch is mediated in part by the response to fatty acid flux (Edwards *et al.,* 2009; Dalebroux *et al.,* 2010). It was shown elsewhere that *in vitro* supplementation of short chain fatty acids (SCFAs), such as propionic acid, will trigger the expression of TP traits (Edwards *et al.,* 2009). Further, the process of differentiation in response to excess SCFAs is mediated by DksA (Dalebroux *et al.,* 2010). PsrA is also associated with differentiation though the precise role remains to be elucidated (Graham *et al*., 2021). The *Pseudomonas* PsrA ortholog is a sensor of long chain fatty acids (LCFAs) of which presence alleviates PsrA-mediated repression of the *fadBA5* β-oxidation operon (Kang *et al*., 2008). To investigate where PsrA also responds to LCFAs, Δ*psrA* strains with and without *in trans* gene expression constructs were grown in BYE broth supplemented with the LCFA palmitic acid to ascertain impact on bacterial growth. In comparison to Lp02, 2-fold growth inhibition of the Δ*psrA* pThy strain was observed which was rescued by *in trans* PsrA expression strongly suggesting that PsrA responds to LCFAs (Figure 11A). Because chromosomal-based DksA expression is present in the Δ*psrA* pThy strain, we next asked if DksA also responded to palmitic acid which could be amplified through *in trans* DksA expression in the Δ*psrA* strain background. While it appears that growth of the Δ*psrA* pVect-*dksA* strain improved in comparison to the Δ*psrA* pThy strain, it was not found to be statistically significant (Figure 11A). Likewise, *in trans* DksA expression in Lp02 did not significantly improve bacterial growth over that of Lp02 alone (Figure 11A). Conversely, *in trans* DksA expression significantly improved bacterial growth over that of Lp02 (Figure 11A). To further investigate the role of DksA, Δ*dksA*-based strains were also assessed. The Δ*dksA* Δ*psrA* pThy strain had 2-fold growth inhibition in comparison to Lp02 (Figure 11B). Growth inhibition of the Δ*dksA* pThy strain was not considered statistically different from that of Lp02 (Figure 11B). Surprisingly, *in trans* PsrA expression in the Δ*dksA* strain could not fully rescue the growth inhibition phenotype. Taken together, these results suggest that while PsrA is largely responsible for the response to LCFAs, DksA also contributes additively to this response lending support to the overlapping roles observed for the two regulatory factors for some of the traits in this study.

**Figure 11:**
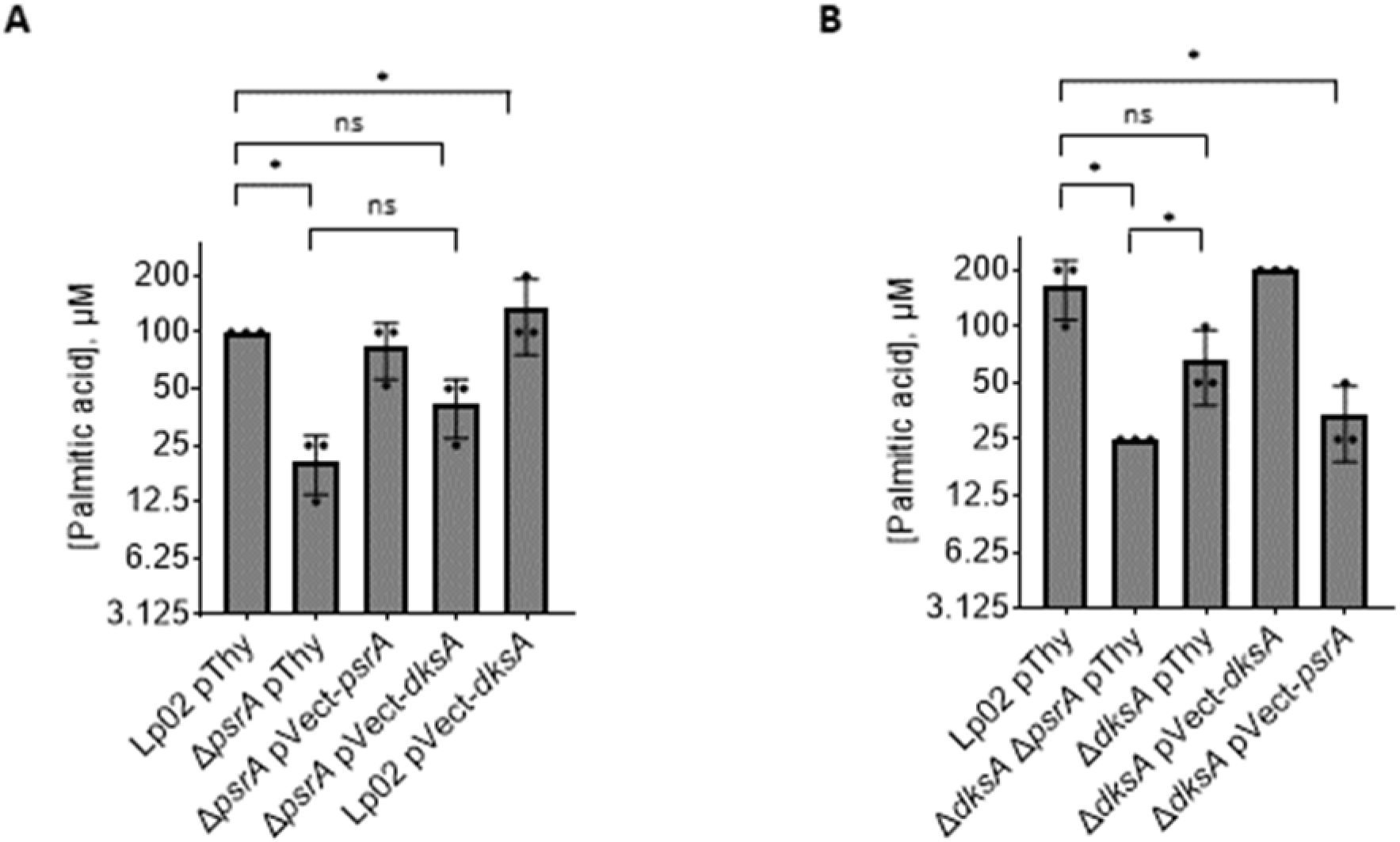
PsrA and DksA influences LCFA-mediated growth inhibition. Parental Lp02 and isogenic strains featuring single or combinatorial genetic mutations (Δ*psrA* and/or Δ*dksA*), with or without *in trans* expression of *psrA* or *dksA* genes, grown with and without palmitic acid supplementation (as 40 mM stock dissolved in ethanol) in BYE media normalized to 0.5% ethanol, in a 96-well microplate for 24 h with shaking at 37°C. Inhibition was defined as growth, as assessed by OD_600_ values, of <50% of that of same strain in a BYE + 0.5% ethanol-only control at the end of 24 h. Bars indicate mean of maximum concentration of PA where indicated strain grew >50% of maximum, over three biological replicates with standard deviation indicated. Dots in bars indicate individual replicate values. Panels A and B were conducted separately. Statistical analyses were done using Student’s unpaired T test: ns = no significance, * = p<0.05.

## Discussion

The biphasic lifecycle of *L. pneumophila* is controlled by the convergence of multiple signals into a sophisticated regulatory network that ultimately acts to induce TP differentiation. The primary signals driving differentiation link to the nutritional status of the environment, mediated largely by induction of the stringent-response and the associated downstream regulatory network in association with RNAP-binding cofactor DksA, alternative sigma factor RpoS, two-component systems and a subset of transcriptional regulators including PsrA (Fonseca and Swanson, 2014; Oliva *et al*., 2018; Graham *et al*., 2024).

This study investigated a spontaneous *dksA1* mutation resulting in a truncated DksA1 protein featuring a deletion within its CC domain (Figure 1) (Perederina *et al,* 2004; Dalebroux *et al.,* 2010). The intermediate phenotype observed in bacteria expressing DksA1 relative to DksA can be explained by comparing their AlphaFold2 models to that of *E. coli* DksA bound to RNAP (Figure 1) (Molodtsov *et al*., 2018). These models predict DksA1 to be physically incapable of coordinating Mg^2+^ in the active site of RNAP due to the truncated loop in the CC domain (Figure 1D, F). However, DksA1 is still predicted to overall form the core CC and GH domains (Parshin *et al*., 2015; Molodtsov *et al*., 2018). This would hypothetically still allow DksA1 to bind into the allosteric binding pocket of RNAP (Figure 1C, S2), thereby recruiting a secondary ppGpp to RNAP. This would partially explain the variance in phenotype of *dksA1* as an intermediary between the WT and knockout strain. Likewise, these structural predictions also fit into the model of the interplay between GreA and DksA as allosteric inhibitors/activators of RNAP competing for the same allosteric site (Parshin *et al*., 2015; Molodtsov *et al*., 2018).

DksA1 (i.e. Δ*dksA1* pThy strain) affected reduced intracellular growth in *A. castellanii* protozoa indicating impaired function that was partially, but not fully, rescued by *in trans* expression of *psrA* (Figure 2A). This surprising finding compelled us to use the DksA1 hypomorph to explore overlapping cellular functions of both regulators, inferring a subset of gene targets jointly regulated by DksA and PsrA. To explore the possibilities that either DksA and PsrA interact with each other, or act in parallel, Δ*dksA* pThy and Δ*dksA* Δ*psrA* pThy strains were subsequently assessed for growth kinetics in *A. castellanii* protozoa (Figure 2B). The Δ*dksA* pThy strain developed a more severely avirulent phenotype than the *dksA1* pThy strain, and was rescued by *in trans* expression of *dksA* or *dksA1* (Figures 2B and 2C), but not by *psrA* (Figure 2B). Notably, the avirulent phenotype of the Δ*dksA* pThy strain contrasts to a previous study showing that a Δ*dksA* insertional mutant strain only exhibited reduced intracellular growth, rather than avirulence, in *A. castellanii* (Dalebroux *et al.,* 2010). While the assays done by Dalebroux *et al*. (2010) and in this study were done in a comparable manner, the discrepancy in findings could be due to differing experimental conditions as the former conducted *A. castellanii* infections at 37°C. Temperature-dependent phenotypes have been seen previously, for example, with Δ*hfq* mutant strains, which only showed an intracellular kinetic defect in *A. castellanii* at 30°C, but not at 37°C (McNealy *et al.,* 2005). We undertook the *A. castellanii* infections at 25°C and showed that the avirulent phenotype of the Δ*dksA* pThy strain was fully complemented by *in trans* DksA expression (Δ*dksA* pVect-*dksA*) indicating that the phenotype was solely due to the genetic deletion of *dksA*. Another possibility for the discrepancy is the possibility of different deletion strategies causing polar effects. Sahr *et al* (2012) annotated *L. pneumophila* Paris strain *dksA* in an operon with an uncharacterized alpha/beta hydrolase *lpp2286* encoded downstream and Lp02-derivative strains are likely similar. However, both our study and that of Dalebroux *et al* (2010) showed the respective Δ*dksA* mutant strains were complementable and thus, the discrepancy is unlikely a polar effect. Additionally, *in trans* expression of DksA1 was sufficient to compensate for lack of DksA indicating that the truncation did not impact essential function of DksA. Lastly, expression of DksA enhanced and rescued intracellular growth in Lp02 and Δ*psrA* strains, respectively. Thus, DksA can compensate for lack of PsrA, and PsrA can partially compensate for the reduced activity of DksA1, but cannot compensate for the absence of DksA intracellularly in *A. castellanii*. Taken together, the interplay of DksA and PsrA in regulating their gene targets is not fully reciprocal, indicating that successful intracellular growth in *A. castellanii* is largely driven by DksA with assistance from PsrA.

The requirement of DksA for intracellular growth is host specific. DksA was shown elsewhere to be dispensable for growth in primary murine macrophages (Dalebroux *et al*., 2010) and for human U937-derived human macrophages as shown in this study (Figure 3). Likewise, the Δ*dksA* Δ*psrA* pThy strain was not impeded for intracellular growth in U937-derived macrophages, a finding similar to that previously found with the Δ*psrA* strain (Graham *et al*., 2021). *In trans* expression of DksA in the Lp02 strain background enhanced intracellular growth in protozoa (Figure 2E), but with little overall impact in U937-derived macrophages (Figure 3B), a trend also seen with overexpression of PsrA (Graham *et al*., 2021). This suggests that elevated levels of either PsrA or DksA on an individual basis imbued a positive effect on replication in *A. castellanii* which does not equate to a substantial phenotype in U937-derived macrophages.

To further delineate the distinct and overlapping roles of DksA and PsrA, *in vitro* approaches were pursued to examine the growth kinetics and phenotypes characteristic of TP-associated traits of *L. pneumophila*. The induction of regulatory mechanisms controlling the transition switch from RP to TP are generally comparably induced *in vivo* (e.g. in protozoa) and *in vitro* (Byrne and Swanson, 1998). In this view, *in vitro* kinetic assays were undertaken to show the interactions between DksA and PsrA were largely growth temperature-independent (Figures 4 and 5). Interestingly, *in trans* expression of DksA1 in the Δ*dksA* strain affected reduced growth rate (Figure 4C) at 37^°^C, but not at 25^°^C (Figure 5D), illustrating that the predicted compromised transcriptional efficiency DksA1 affects growth rates at higher temperatures. This is further supported by the growth profile of Δ*dksA* pVect-*dksA1* which resembled that of Lp02 in *A. castellanii* (Figure 2C). The slow *in vitro* growth phenotype did not apply to the Lp02 pVect-*dksA1* strain bearing chromosomally-expressed DksA, inferring that *dksA1* is not a dominant-negative allele (Figure 4F). The growth profiles of *in vitro* grown *dksA1* pThy and *dksA1* pVect-*psrA* were similar to each other at both at 25^°^C (Figures 4A, 4B) and 37^°^C (Figures 5A, 5B), but not in protozoa where a two-log_10_ recovery was achieved through *in trans* expression of PsrA inferring shared gene targets.

Other TP-associated phenotypes include cellular filamentation, culturability, and pigmentation are mediated by DksA independently as previously reported (Dalebroux *et al.,* 2010). Lack of DksA resulted in unusual colony morphology (Figure 6) and filamentation, particularly in stationary phase (Figures 7-8). Recently, DksA was ascertained to have a pivotal role in cell division in *E. coli*. Binding of ppGpp to DksA/RNAP indirectly activates the divisome complex component FtsZ to modulate cell length and division (Anderson *et al*., 2023). The absence of either DksA or ppGpp promotes cell filamentation for different reasons. Lack of DksA prevents activation of cell division by ppGpp resulting whereas lack of ppGpp causes DksA to switch from an activator to an inhibitor of FtsZ. Thus, by orthologous function, it is likely that DksA behaves in the same manner in *L. pneumophila*. The dramatically reduced bacterial culturability of Δ*dksA* pThy strain (Figure 9) demonstrates the inability of the strain to physiologically respond to nutrient deprivation and other stressors, which is consistent with the importance of DksA to the stringent response. Biologically, PsrA does not contribute to this phenotype as demonstrated by transcriptomic analyses that showed few stress-related genes under PsrA regulation (Graham *et al*., 2021). Because variability in filamentation and culturability affected by lack of DksA can impact optical density to titre ratios of stationary phase bacteria, extreme care was taken in this study to ensure that Δ*dksA*-based strains were recovered from ≤ 3-day-old BCYE-agar plates to minimize this variability for *in vitro* growth curves and inoculum titre for *in vivo* kinetics. Lastly, pigmentation comprises a brown tyrosine-based metabolite produced post-exponentially by *L. pneumophila* as an adaptation to environmental stress, potentially serving for photoprotection, iron acquisition, or as a biocide to enhance *L. pneumophila* competitiveness in biofilms (Steinert *et al.,* 1995; Chatfield and Cianciotto, 2017; Levin *et al.,* 2019). As reported elsewhere, DksA is a major driver of pigment production (Figure 10) (Dalebroux *et al.,* 2010). Thus, these traits, other than cell length, do not seem measurably impacted by the cellular levels of PsrA, indicating that PsrA may impact virulence specific pathways rather than cellular physiology.

Another phenotypic trait associated with TP differentiation is fatty acid flux which is linked through SpoT-mediated stringent induction, with either DksA or ppGpp, sufficient to arrest growth upon supplementation with SCFA (Edwards *et al.,* 2009; Dalebroux *et al.,* 2010). Here, we show that like its ortholog in *Pseudomonas* spp., PsrA is responsive to fatty acids suggesting a conserved functional role in sensing LCFA in the environment alongside SpoT. Further, both DksA and PsrA must be present to ensure a proper LCFA response (Figure 11) (Kang *et al*., 2008). However, the precise mechanistic details of the PsrA-mediated response remain undefined warranting further investigations.

In summary, this study explored the phenotypic roles of spontaneously mutated DksA1 and PsrA, and uncovered a synergistic genetic relationship between PsrA and DksA in terms of virulence regulation. The bacterial response to LCFA suggests a novel regulatory link to the stringent response pathway, further solidifying the mechanistic importance of both DksA and PsrA as contributing regulators of the biphasic lifestyle of *L. pneumophila.* Because only the predicted, rather than the functional, basis of the *dksA1* phenotype has been derived, these results cannot distinguish the distinct effects of ppGpp recruitment and competitive sequestration of RNAP from GreA. With these limitations in mind, our findings suggest that PsrA seems to redundantly act as a bypass for DksA in virulence but has little impact on other processes, suggesting a more targeted mechanism. The overlapping phenotypes presented in this study are consistent with the two distinct mechanisms by which DksA and PsrA act, and thus the following model is proposed based on a scenario in which a promoter is repressed by either DksA or PsrA (Figure 12). Absence of DksA would permit GreA to act as the sole secondary channel binding protein, thereby allowing increased efficiency in transcription initiation, and should there be elevated levels of PsrA, it would result in increased promoter blockage providing compensatory repression. Absence of PsrA would leave the promoter more accessible for RNAP, but this could be counteracted should there be elevated levels of DksA resulting in reduced transcriptional efficiency. Alternatively, a promoter not subject to PsrA-mediated influence would be exempt from the compensatory effects, and thus be wholly dependent on the dynamics between GreA and DksA. Nevertheless, the proposed co-regulator duality could explain why transcriptomic analyses did not overtly reveal a large number of PsrA-regulated targets (Graham *et al*., 2021).

**Figure 12:**
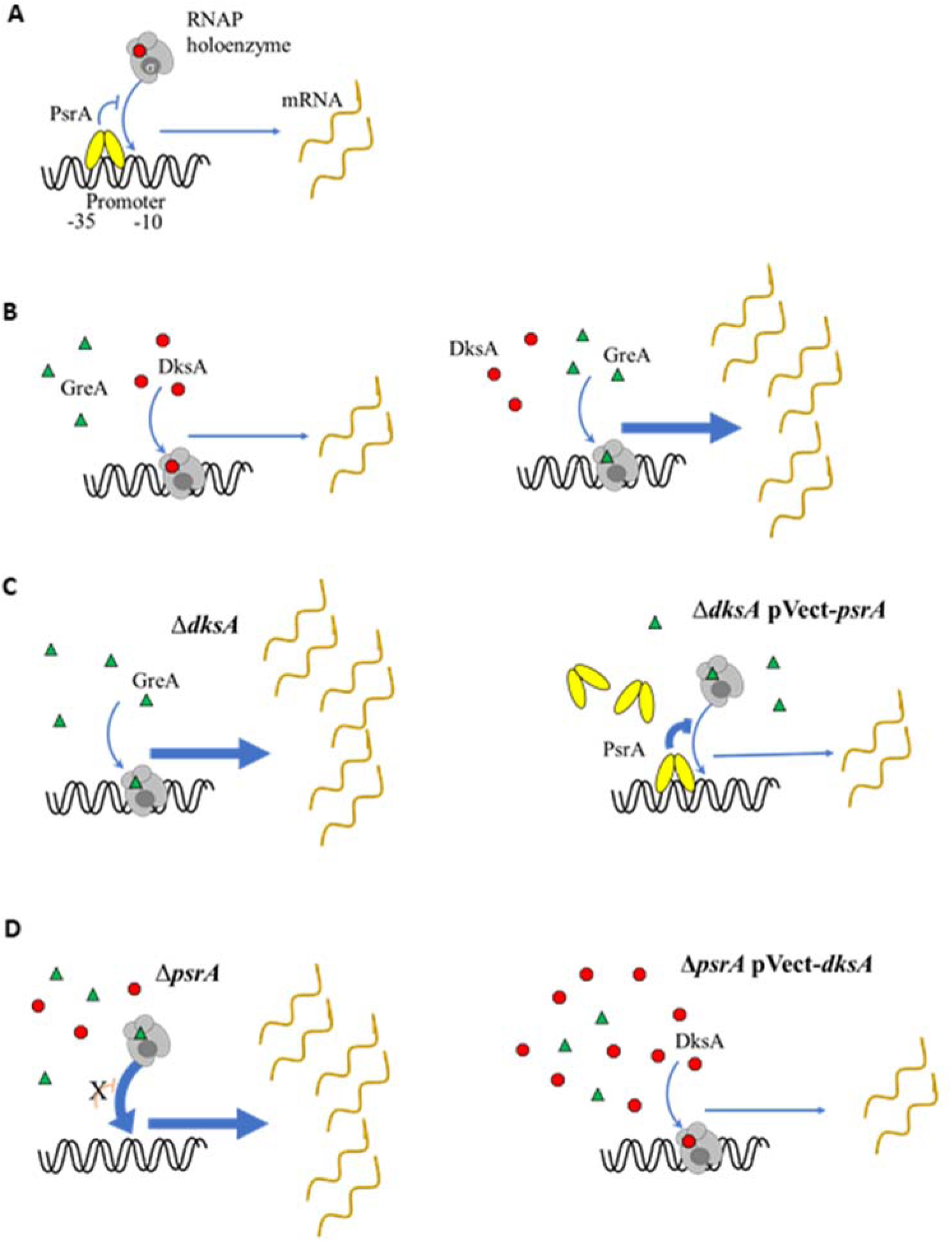
Model of independent mechanisms of DksA and PsrA. A hypothetical promoter is repressed by both PsrA and DksA (A) RNA polymerase (RNAP) holoenzyme is targeted to -10 and -35 boxes by a sigma factor. PsrA (oval dimers) binds a defined site in the promoter and inhibits RNAP recruitment. (B) RNAP binds to GreA (triangles) or DksA (filled octagons) interchangeably, with enhanced transcription initiation rates occurring while bound to GreA versus DksA. (C) In a Δ*dksA* strain, only GreA is available for RNAP thus elevating transcription levels, but *in trans psrA* overexpression generates higher levels of PsrA, strengthening inhibition of RNAP recruitment to compensate. Conversely (D) the Δ*psrA* strain allows uninhibited RNAP-promoter binding, increasing transcription, but this is offset by overexpressing *dksA in trans*, increasing the proportion of RNAP bound to transcription- inhibiting DksA thereby generating the compensatory effect.

Thus, future transcriptomic studies will include the Δ*dksA* Δ*psrA* strain to compare against data obtained for individual Δ*psrA* and Δ*dksA* strains to discern distinct and overlapping roles. These studies could also be expanded to include the ppGpp^0^ (Δ*relA* Δ*spoT*) strain to further explore the interface with PsrA and ppGpp in relation to the link between PsrA and the fatty acid response.

## Acknowledgements

We thank M. Swanson (University of Michigan) for the kind gift of pBH6119 and Lp02, J. Vogel (Washington University) for the kind gift of pSR47s and pJB908, R. R. Isberg (Tufts University) for the kind gift of Lp03. This work was supported by Natural Sciences and Engineering Research Council (NSERC) Discovery Grant funds awarded to A.K.C.B (RGPIN- 2019-05490), T.dK. (RPIN-2017-04970) and G.P. (RGPIN-2018-04968), and Canadian Foundation for Innovation (CFI) Grant funds to A.K.C.B. Research personnel were supported by University of Manitoba Faculty of Graduate Studies (C.G., T.M, T.P., J.B.), Research Manitoba Master’s Studentship Award (T.P.), and NSERC CGS Masters (T.M, A.G.) and Doctoral (T.M., T.P, A.G.).

