## Supplemental information for "Transcription Factors DksA and PsrA are synergistic contributors to *L. pneumophila* virulence in *Acanthamoeba castellanii* protozoa"

List of supplementary information:

Figures S1 to S2

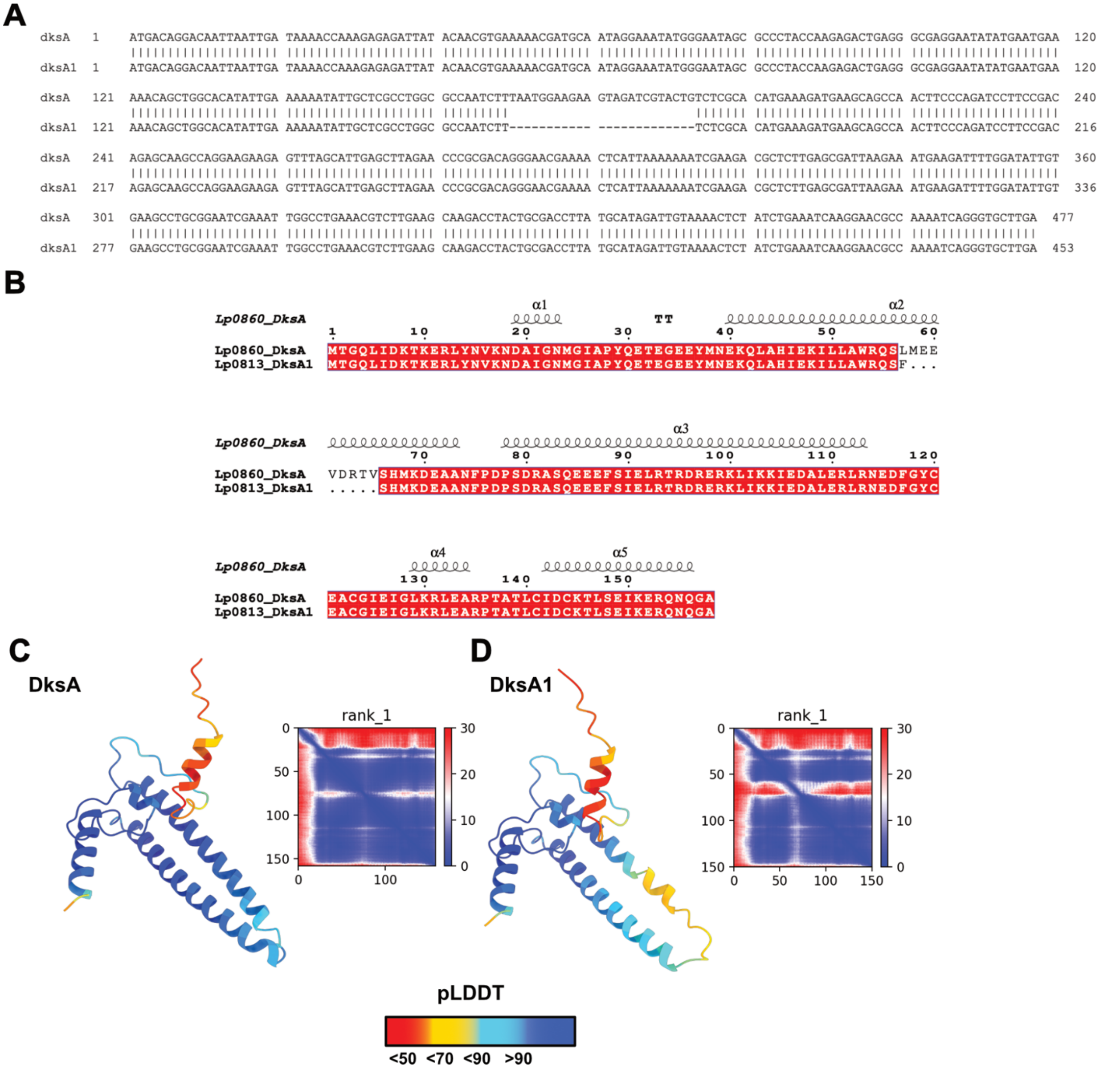


**Figure S1: The *dksA1* allele encodes a truncated protein.** Alignment of (A) DNA sequences of parental Lp02 *dksA* (from Rao *et al*., 2013) and spontaneous mutant allele *dksA1*; and (B) amino acid sequences of Lp02 DksA and DksA1 as encoded by genes in (A). Protein sequences were aligned using ClustalOmega (https://www.ebi.ac.uk/jdispatcher/msa/clustalo) and visualized by Espript3 (https://espript.ibcp.fr/). Ascribed secondary structure was generated using the AlphaFold model of DksA in (C). AlphaFold2 models of DksA (C) and DksA1 (D) are shown and colored by pLDDT (prediction confidence). In lay plots show the exact pLDDT by residue.

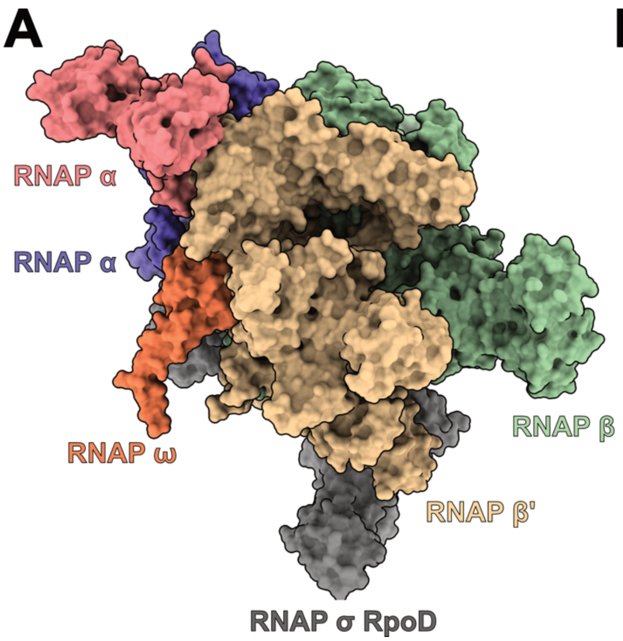


**Figure S2: Structural overlay of Lp02 DksA and DksA1 with an *E. coli* DksA:RNAP complex** Comparison of RNAP (PDB ID: 5VSW) modelled without (A) and with DksA (B) in the allosteric site. DksA (5VSW) is modelled in beige, while Lp02 DksA and DksA1 are modelled in blue and salmon respectively. RNAP subunits are labeled and colored. When viewed with matchmaker (UCSF Chimera) the secondary structure of the coiled-coiled domains are nearly identical, and allow the formation of the conserved secondary ppGpp binding site.
